# Integrated characterization of cell types, states and molecular programs in disseminated appendiceal neoplasms

**DOI:** 10.1101/2023.09.28.559817

**Authors:** Linh T. Bui, Xu Cao, Jinhui Wang, Fan Meng, Mingye Feng, Leonidas Arvanitis, Rifat Mannan, Yanghee Woo, Kamran Idrees, Nicholas E. Banovich, Mustafa Raoof

## Abstract

Appendiceal neoplasms include a heterogeneous group of epithelial and non-epithelial tumors with varying malignant potential. Despite the rise in incidence of appendiceal neoplasms in recent years, limited progress has been made in the understanding, management and therapeutic treatment. To comprehensively characterize the cell types and molecular mechanisms driving cellular remodeling in epithelial appendiceal neoplasms, we performed an integrated scRNA-seq study. We analyzed 126,998 cells from 16 appendix samples (11 peritoneal metastases samples, 5 healthy controls) and identified 33 distinct cell types/cell states with seven being cancer-specific. Highlights of our study include the characterization of tumor cells across the histologic spectrum, the identification of a novel cancer-associated-fibroblast (CAF) subtypes (fiCAFs) and the identification of pathologic-specific cellular crosstalk between tumor cells and the tumor microenvironment (TME). Together, our study provides a high-resolution insight into the complexity and heterogeneity of epithelial appendiceal neoplasms and a valuable resource for therapeutic strategies.

## Introduction

Appendiceal neoplasms are rare accounting for less than 1% of all gastrointestinal cancers^1^. In the US, approximately 1500 cases will be diagnosed each year^2^. Between 2000 and 2016, the age-adjusted incidence of malignant appendiceal neoplasms increased by 232%^1^. Despite the rising incidence, there has been limited progress in the understanding and management of appendiceal neoplasms.

Broadly, appendiceal neoplasms are classified as either epithelial (arising from epithelial cells) or non-epithelial (arising from neuroendocrine cells)^3^. Among epithelial appendiceal neoplasms, there is a strong propensity for peritoneal metastases and a marked heterogeneity in clinical outcomes; with long-term survival for metastatic disease ranging from 10% - 90%^2–4^. Histologic classification of epithelial appendiceal neoplasms is the mainstay of clinical decision making but achieving a consensus histologic classification scheme has been a significant challenge in the field^5–8^. To adequately capture the histologic heterogeneity, several classification systems have been proposed in the past^5–8^. According to the most recent PSOGI consensus^8^ and WHO classification schemes^9^, epithelial appendiceal neoplasms can be classified as: mucinous appendiceal neoplasm/ adenocarcinoma; goblet cell adenocarcinoma (GCA); or non-mucinous (colonic-type) adenocarcinoma. Within each histologic subtype, grade is a strong prognostic factor, and the distribution of grade varies by histologic type^1^. For instance, most individuals with peritoneal dissemination from mucinous neoplasms/ adenocarcinomas have a low-grade histology and have 10-year survival greater than 80% after complete cytoreduction^10^; whereas the majority of GCAs and non-mucinous adenocarcinomas with peritoneal metastases have a high-grade histology and 10-year survivors despite complete cytoreduction are rare^11–13^. Within existing histologic classification schemes there remains a remarkable heterogeneity in response to treatment, and clinical outcomes; necessitating the development of improved predictive and prognostic methods.

Several groups have explored the utility of integrating histologic findings with somatic mutation profiling for better characterization of appendiceal neoplasms^14–17^. These studies have demonstrated distinct genomic alterations that correlate with histologic subtypes of appendiceal neoplasms. For instance, mucinous neoplasms/ adenocarcinomas have activating mutations in GNAS (52%) and KRAS (72%) but TP53 alterations are less common (26%). In contrast, activating GNAS mutations are uncommon (15%) in non-mucinous (colonic-type) adenocarcinomas but KRAS mutations (70%) and TP53 mutations (55%) are more common. GCAs have a unique mutation profile with TP53 mutation being the most common alteration occurring in 26% of patients, among other less common mutations (e.g., KRAS, SMAD4, SOX9). An all-encompassing molecular classification scheme of appendiceal epithelial neoplasms has been proposed with four categories: KRAS-predominant (KRAS-mutant, GNAS-and TP53-wildtype); GNAS-predominant (GNAS-mutant, with or without KRAS mutation, and TP53-wildtype); TP53-predominant (Any TP53-mutant); or triple negative^14^. While this classification has prognostic utility, models that included histologic classification with molecular features performed the best (AUC 0.75) which was a marginal improvement over histologic classification based on grade alone (AUC 0.68).

Studies using bulk gene expression analyses on tumors from patients who have undergone complete cytoreduction have delineated two prognostic sub-categories among low-grade mucinous adenocarcinomas/ neoplasms (a high-risk and a low-risk subtype)^18^. Further work focusing on transcriptomic subtyping using Nanostring platform demonstrated that a 17-gene signature (7 – immune genes, and 10 – oncogenes) could stratify mucinous neoplasms/ adenocarcinomas into three molecular sub-types: immune-enriched, oncogene-enriched, and mixed^19,20^. Immune-enriched tumors demonstrated the best prognosis while oncogene-enriched tumors demonstrated the worst prognosis after cytoreduction.

Despite these advances in molecular characterization and identification of unique driver mutations (GNAS and KRAS); there has been little therapeutic advancement as these mutations have not been druggable to date. Early-stage cancers can be treated definitively with surgery, and selected patients with metastatic disease derive long-term benefits from cytoreductive surgery and hyperthermic intraperitoneal chemotherapy (HIPEC). For unresectable disease, however, there are no proven therapies. Even though there is strong evidence that appendiceal neoplasms are histologically and molecularly distinct from colorectal cancers (CRC), combination chemotherapy that is used for CRC is often implemented for appendiceal cancers with mixed results^15–18^. Furthermore, appendiceal neoplasms are considered immunologically cold and have not been reported to respond to currently used immune checkpoint-blocking therapies^17^.

To identify novel therapeutic vulnerabilities in appendiceal neoplasms and improve patient outcomes, there is an urgent need for a deeper molecular understanding of the phenotypic diversity of cancer and non-cancer cell populations across a spectrum of histologic subtypes. We hypothesized that defined histologic subtypes will not only have unique epithelial molecular programs and cell states; but also unique tumor microenvironments with heterogeneous non-cancer cell populations. Here, we use comprehensive single-cell RNA sequencing to reveal the global landscape of disseminated appendiceal neoplasms and uncover molecular pathways that may be leveraged for improved risk stratification or exploited for therapeutic development.

## Results

### Integrated scRNA-seq analyses of appendiceal cancer

To characterize the cell types and molecular mechanisms driving cellular remodeling in epithelial appendiceal neoplasms, we performed an integrated scRNA-seq study on 126,998 cells from 16 patient samples (11 peritoneal metastases and 5 normal appendix samples) (Fig. 1a). The 11 peritoneal metastases samples are further classified based on histology into four pathology groups: low-grade mucinous adenocarcinoma (LGMA, n = 4), low-grade appendiceal mucinous neoplasm (LAMN, n = 2), moderate/high non-mucinous adenocarcinoma (MHNA, n = 3) and goblet-cell adenocarcinoma (GCA, n = 2) (Fig. 1b, Supplementary Fig. 1, Supplementary Table 1). Mutation profiling assays using a comprehensive genomic profiling assay (GEM ExTra) confirmed previous studies regarding different mutation patterns in appendiceal pathologies (Fig. 1f, Supplementary Table 2). Using marker-based annotations, we identified eight cell lineages (Fig. 1c-f, Supplementary Fig. 2) that were further characterized into 33 distinct cell types/cell states with seven being cancer-specific (Supplementary Fig. 3).

**Fig. 1:**
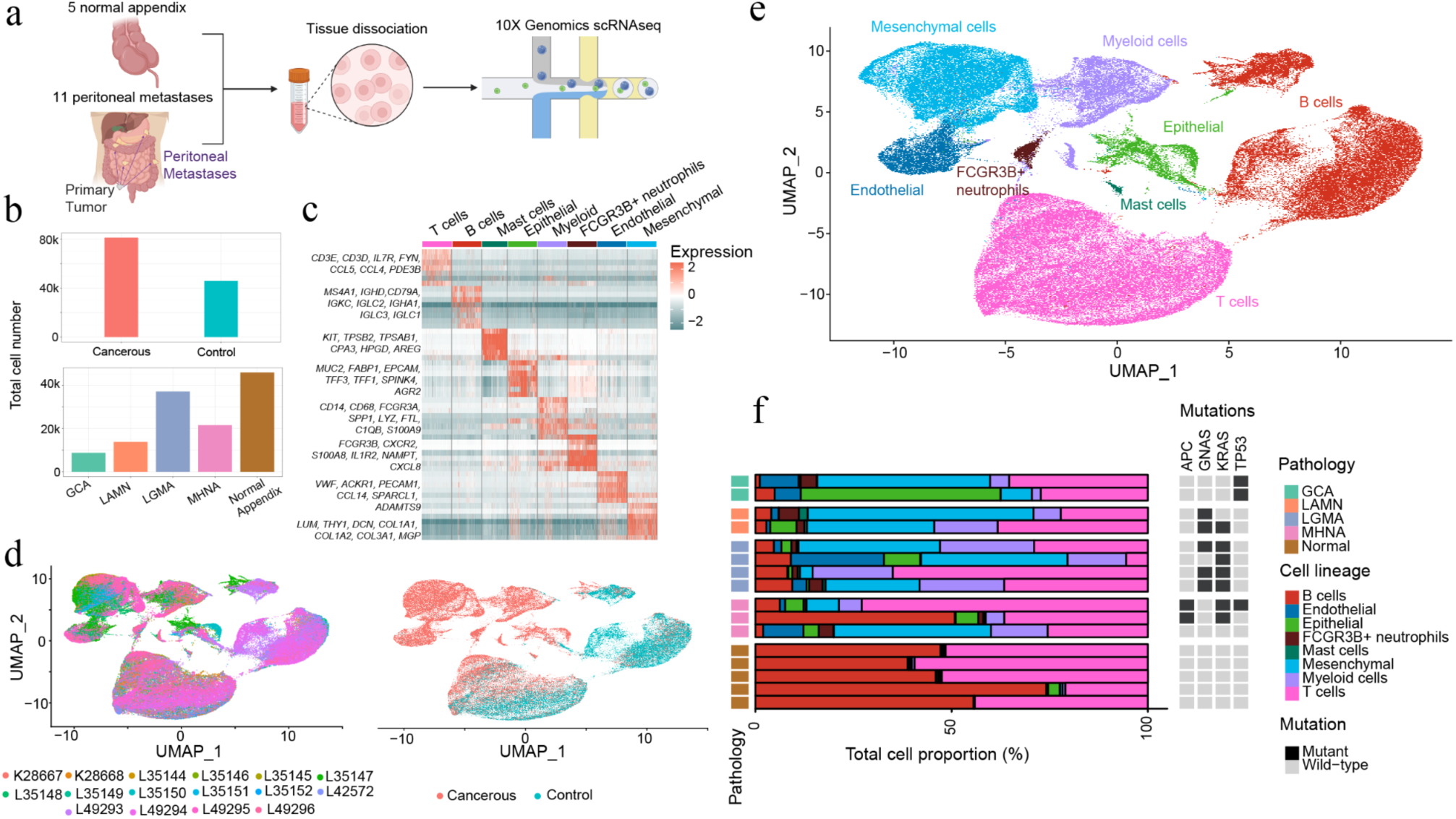
Integrated scRNA-seq analyses of appendiceal cancer. **a** Schematic illustration of the scRNA-seq experiment in the study. **b** Total number of cells broken down in terms of cancerous vs. control samples or of different pathology groups in the study. **c** Heatmap depicts cell lineage-specific marker genes used to annotate the different cell lineages. **d, e** Dimensional reduction view of 126,998 cells in the study, by sample ID and by status (**d**) or by cell lineages (**e**). **f** Cell type proportion of different cell lineages per sample, with mutation profile of the individual sample. Mutations were detected using the genomic profiling test GEM ExTra^21^.

### Characterization of epithelial cells in appendiceal neoplasms

The epithelial cell lineage in our dataset consists of 5,205 individual cells, with most cells derived from cancerous samples (4,763 cells - 91.5% total epithelial cells). Unsupervised clustering grouped the majority of tumor cells into pathology-specific clusters (Fig. 2a), demonstrating a high degree of transcriptional divergence across the histologic spectrum (Supplementary Fig. 6a-b). Importantly, peritoneal metastases from LGMA and LAMN many times are histologically indistinguishable without looking at the primary tumors. However, our single-cell analysis demonstrated distinct clusters for the two entities. Compared to normal appendix cells, tumor epithelial cells contain multiple copy number variants (CNVs), with CNVs being either pan-appendiceal or more unique to each subtype (Fig. 2a-b, Supplementary Fig. 4). Among the four pathologies, epithelial cells derived from GCA samples exhibit more CNVs such as duplication chr9, chr19 and chr20, and a subset of these cells have duplication in chr16.

**Fig. 2:**
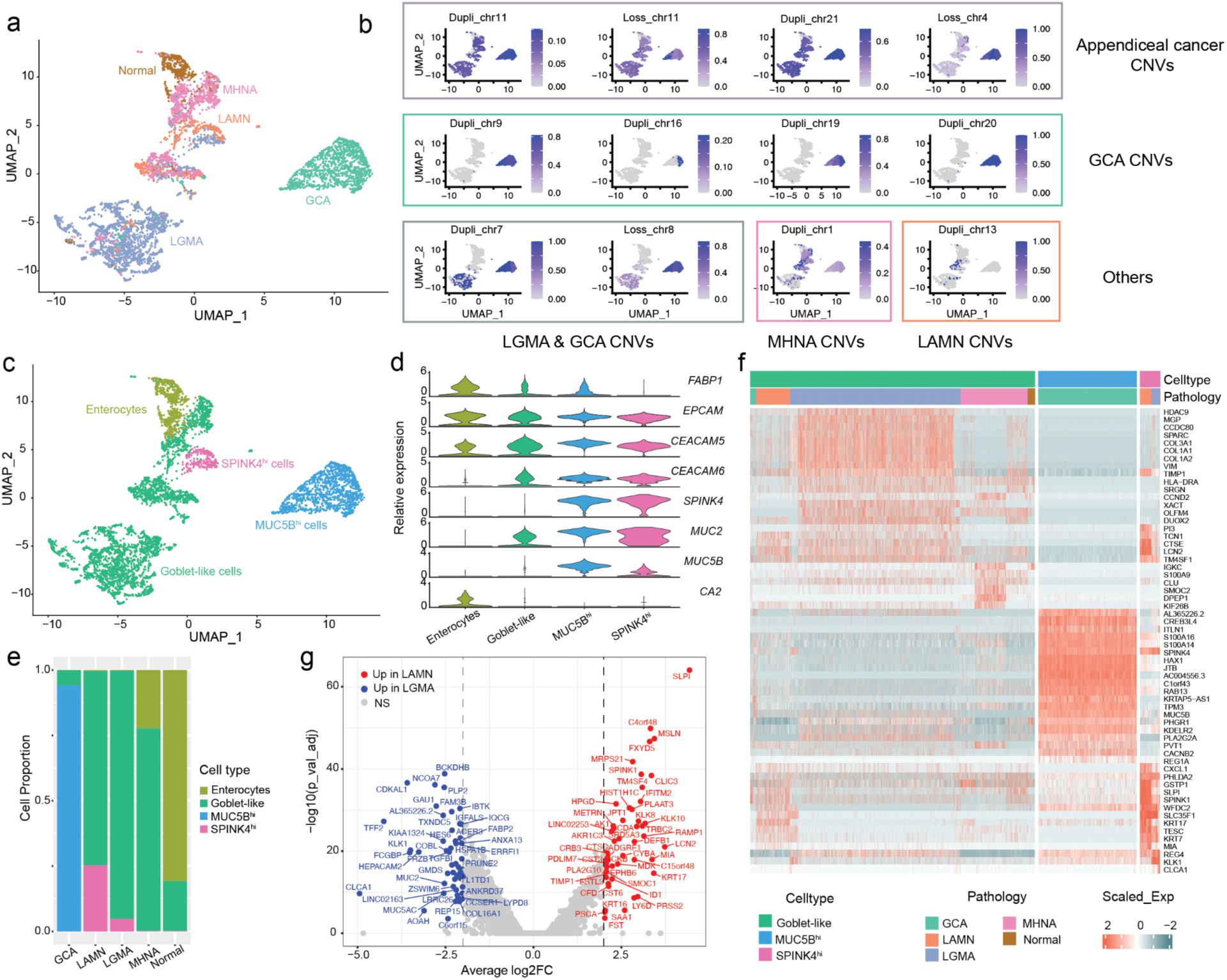
Epithelial cell characterization in appendiceal cancer. **a** Unsupervised clustering of the epithelial cells results in cells grouping in different pathology groups. **b** CNV profiles of pan-subtype appendiceal cancer or subtype (GCA, LGMA, MHNA, or LAMN). Plots display the proportion of expressed genes within the CNVs identified by infercnv^29^. **c** UMAP plot displays epithelial cells grouping by cell types. **d** Violin plots depict the relative expression of epithelial marker genes in each cell type. **e** Epithelial cell type proportion distribution in each pathology group. **f** Heatmap displays scaled expression levels (normalized and scaled z-scores) of the top highly expressed genes in the goblet-like cells, *MUC5B*^hi^ cells and *SPINK4*^hi^ cells split by pathology groups. **g** Volcano plot depicts differentially expressed genes (DEGs) in LAMN vs. LGMA *SPINK4*^hi^ cells; red: significant upregulated (FDR ≤ 0.05, avg_log2FC ≥ 2) in LAMN *SPINK4^hi^* cells, blue: significant upregulated in LGAM *SPINK4^hi^* cells (FDR ≤ 0.05, avg_log2FC ≤ -2), NS: not significant.

Using marker-based annotations, we identified four major cell types in the epithelial cell lineage including enterocytes, goblet-like cells, *SPINK4*^hi^ cells and *MUC5B*^hi^ cells (Fig. 2c-d, Supplementary Fig. 5a-b). Enterocytes are the dominant cell types in normal epithelial cells and express a high level of carbonic anhydrase 2 (*CA2*). The majority of epithelial cells derived from cancerous samples (goblet-like cells, *MUC5B*^hi^ cells, and *SPINK4*^hi^ cells) have goblet cell characteristics (*MUC2+, REG4+* or *TFF3+*) and express high levels of the tumor marker cell adhesion molecules 5 and 6 (*CEACAM5, CEACAM6*) (Fig. 2d, Supplementary Fig. 5c). *MUC5B*^hi^ cells additionally express high levels of mucin genes (*MUC5B* specifically) besides goblet cell markers (*MUC2, TFF3*); and are the predominant cancer cell population in GCAs. In addition, a subset of goblet-like cells –denoted as *SPINK4*^hi^ cells– express high levels of *SPINK4* but low levels of *MUC5B*. *SPINK4* is a favorable prognostic factor in CRC^22^. *SPINK4*^hi^ cells were exclusively found in LAMN and LGMA samples; both of which have a more favorable prognosis than MHNA. Expectedly, *SPINK4*^hi^ cells were more prevalent in the prognostically more favorable LAMN compared to LGMA (Fig. 2e). Interestingly, unlike the other three mucinous pathology grades, goblet-like cells derived from MHNA have elevated expression of *TFF3* but not *MUC2* (Fig. 2d, Supplementary Fig. 5). Lack of *MUC2* expression in CRC is associated with poor prognosis^23^, consistent with the worst outcomes of MHNA compared to LGMA and LAMN. The variation in the expression of goblet cell markers in epithelial cells across the histologic spectrum indicates distinct epithelial lineages with differences in the molecular mechanisms driving tumor development.

To further characterize the molecular differences among appendiceal pathologies, we study gene expression changes in the epithelial cells derived from the four histology groups. Goblet-like cells derived from different pathology groups display differences in gene expression profile (Fig. 2f). We observed inter-patient heterogeneity in goblet-like cells derived from a patient (L35147) with distinct gene expression profile compared to other samples in the LGMA group (Fig. 2f, Supplementary Fig. 6). Interestingly, these cells express fibroblast markers such as *COL1A1*, *COL1A2*, *COL3A1* and *VIM*, suggesting an epithelial-mesenchymal transition (EMT) state. Compared to goblet-like cells, *MUC5B^hi^* cells express high levels of genes involved in tumor formation and progression such as Interlectin-1 (*ITLN1*)^24^, cAMP responsive element binding (*CREB3L4*)^25^, HCLS1-associated protein X-1 (*HAX1*)^26^, and a member of the RAS oncogene family (*RAB13*)^27^. To further characterize the differences between LAMN and LGMA *SPINK4^hi^*cells, we performed a differential expression (DE) analysis (Fig. 2g). LAMN *SPINK4^hi^* cells express high levels of *SLPI* (a secretory leukocyte protease inhibitor produced by healthy intestinal epithelial cells and has a promoting role in tumor metastasis in CRC^28^), the interferon *IFITM2* and the mesothelial cell marker *MSLN*. On the other hand, the trefoil factor 2 (*TFF2*), annexin A13 (*ANXA13*) and the lncRNA *GAU1* are among the genes being upregulated in the LGMA *SPINK4^hi^* cells. We also performed a DE analysis in non-*SPINK4*^hi^ goblet-like cells from LGMA and LAMN samples. This demonstrated upregulation of 600 genes (FDR ≤ 0.05 & avg_log2FC ≥ 1) in LAMN samples, whereas 6 genes (FDR ≤ 0.05 & avg_log2FC ≤ -1) were upregulated in LGMA samples (Supplementary Fig. 6c, Supplementary Table 5).

### Cellular remodeling in the tumor immune cell lineage

Despite being an important factor in the development of cancer, the appendiceal tumor immune microenvironment is not well understood. To identify changes in the immune cell lineage between cancerous and normal cells, as well as among different pathologies, we explored the immune compartment of the scRNA-seq dataset. We analyzed 76,365 lymphocytes and identified nine distinct cell types, five T cell and four B cell subtypes (Fig. 3a, Supplementary Fig. 7). Normal lymphocytes comprise mostly B cells, and there is a significant reduction in B lymphocytes in appendiceal tumor tissues. On the contrary, T-cell subtypes (CD4+ T cells, CD8+ T cells and NK cells) had a higher distribution in tumor tissues (Fig. 3b). While CD4+ T cells were similarly distributed across histology, CD8+ T cell abundance varied with highest prevalence in LGMA and LAMN samples, intermediate in GCA samples, and low in MHNA samples. Interestingly, plasma B cells were predominantly present in LGMA samples compared to other cancer histology.

**Fig. 3:**
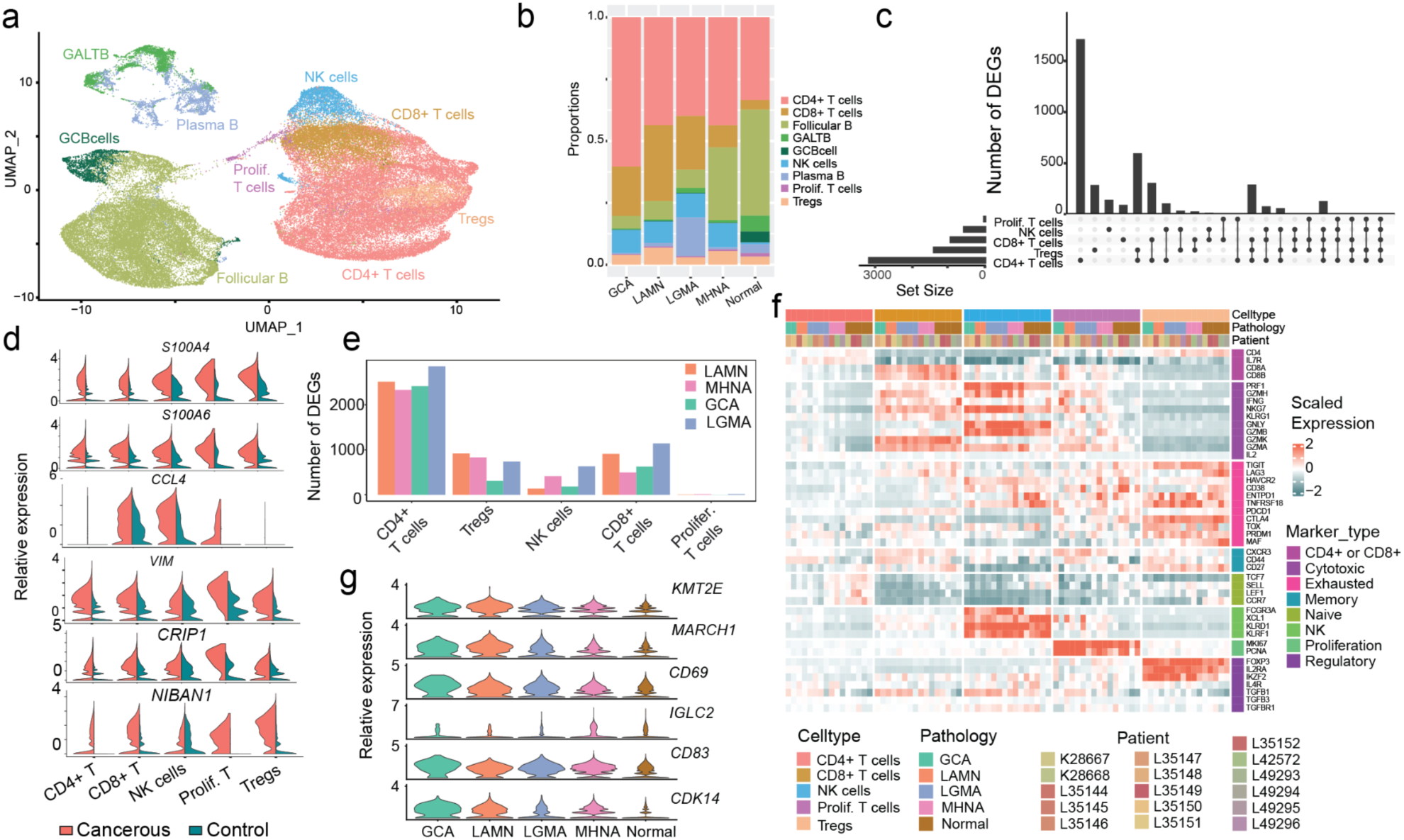
Adaptive immune cell profiles in appendiceal cancer. **a** UMAP displays the nine cell types identified in B and T lymphocytes in both cancerous and control samples; GALTB: gut-associated lymphoid tissue B cells, GCBcells: germinal center B cells. **b** Proportion of B and T cells in each pathology group. **c** Upset plot displays shared and unique DEGs in the comparison between cancerous and control cells for each cell type in T lymphocytes. **d** Upregulation of cancer progression-promoting genes in different T lymphocytes from cancerous compared to healthy tissues. **e** Numbers of DEGs between different pathology vs. control in each T cell type. **f** Heatmap depicts the average relative expression (normalized and scaled z-score) of different T-cell marker genes in each T-cell cluster per pathology group. **g** Expression of some immune response marker genes in the follicular B cells across pathologies.

To study the differences in gene expression of T lymphocytes derived from cancerous and normal tissues, we performed a DE analysis (Fig. 3c). We observed an upregulation of cancer progression-promoting genes (*S100A4*, *S100A6, CCL4*)^30–32^, apoptosis regulator (*NIBAN1*)^33^, EMT marker (*VIM*), and CRC progression (*CRIP1*)^34^ in the cancerous T lymphocytes compared to healthy control (Fig. 3d). While vimentin (*VIM*) is known as an EMT marker^35^, recent research reported high expression of *VIM* in lymphocytes isolated from septic shock and sepsis patients^36^. In addition, we characterized pathology-specific changes in gene expression in T-cell subtypes and reported a collective set of DEGs between each pathology and control for each T cell subtypes (Fig. 3e, Supplementary Fig. 7b, Supplementary Fig. 8, Supplementary Table S5).

Among T lymphocytes, cancerous CD8+ T cells and NK cells exhibit a cytotoxic phenotype compared to control cells, while expression of exhaustion markers like *ENTPD1, TNFRSF18* is elevated in cancerous Tregs (Fig. 3f). In agreement with current literature on appendiceal neoplasms being immunologically cold^17^, key immune checkpoints such as *PD1 (PDCD1)*, *PD-L1 (CD274), PD-L2 (PDCD1LG2*), *LAG3, TIGIT* and *CTLA4* are expressed at a low level in all cells (Supplementary Fig. 9). However, we detected dysregulation of other immune checkpoint genes in the cancerous cells. For example, many members of human leukocyte antigens (HLA) such as *HLA-DRA, HLA-DRB5, HLA-DRB1, HLA-DQA1, HLA-DQB1, HLA-DPA1, HLA-DPB1* and members of the tumor necrosis factor receptor superfamily such as *TNFRSF4, TNFRSF9, TNFRSF14, TNFRSF18* are upregulated in cancerous Tregs and proliferating T cells. Certain members of the killer cell immunoglobulin-like receptors (*KIR2DL1, KIR2DL3, KIR3DL1* and *KIR3DL2*), key regulators of NK cell function, are upregulated in the cancerous NK cells compared to control; with the exception of *KIR2DL4* which is higher in NK cells from normal appendix (Supplementary Fig. 9).

Of all B-cell subtypes, follicular B is the only cell type present with an adequate number in all histology groups. We performed a DE analysis in follicular B cells between each pathology and control to identify pathology-specific transcriptional changes in follicular B cells (Fig. 3g, Supplementary Fig. 8). Cancerous follicular B cells express high levels of cancer-associated genes such as the *KMT2E^37^* (a member of the histone methyltransferase family), *MARCH1^38^*, *CD69^39^* (an anti-tumor immunity regulator), immunoglobulin *IGLC2^40^*, MAPK signaling interactor *CD83^41^* and the cyclin-dependent kinase CDK14^42^.

To define the major population and subpopulation of the tumor-infiltrating myeloid cells, we sub-clustered and analyzed the myeloid cell lineage. Using marker-based annotation, we classified the myeloid cell clusters into seven cell types including three dendritic cell populations (pDCs, cDC1 and cDC2), two monocyte populations (monocyte-like and C1Qhi monocytes) and two macrophage populations (macrophages and *SPP1*+ macrophages) (Fig. 4a-c, Supplementary Fig. 10). C1Qhi monocytes were characterized by high expressions of complement 1q chains (*C1QA*, *C1QB*, *C1QC*), monocyte markers (*FCGR3A, CD14*). Additionally, C1Qhi monocytes express macrophage markers (*LYVE1*, *NLRP3, PLTP*) and a low level of secreted phosphoprotein 1 (*SPP1*). Monocyte-like cells express high levels of *FCN1* and S100A family genes (*S100A8*, *S100A9*) and *SPP1*+ macrophages express high levels of *SPP1*. Moreover, apolipoprotein E (*APOE*), a gene associated with an anti-inflammatory phenotype^43^, is upregulated in C1Qhi monocytes and *SPP1+* macrophages (Fig. 4b). We observed a variation of myeloid cell distribution across pathologies, with MHNA samples having an increase in *SPP1*+ macrophages and monocyte-like but fewer C1Qhi monocytes than other pathologies (Fig. 4c).

**Fig. 4:**
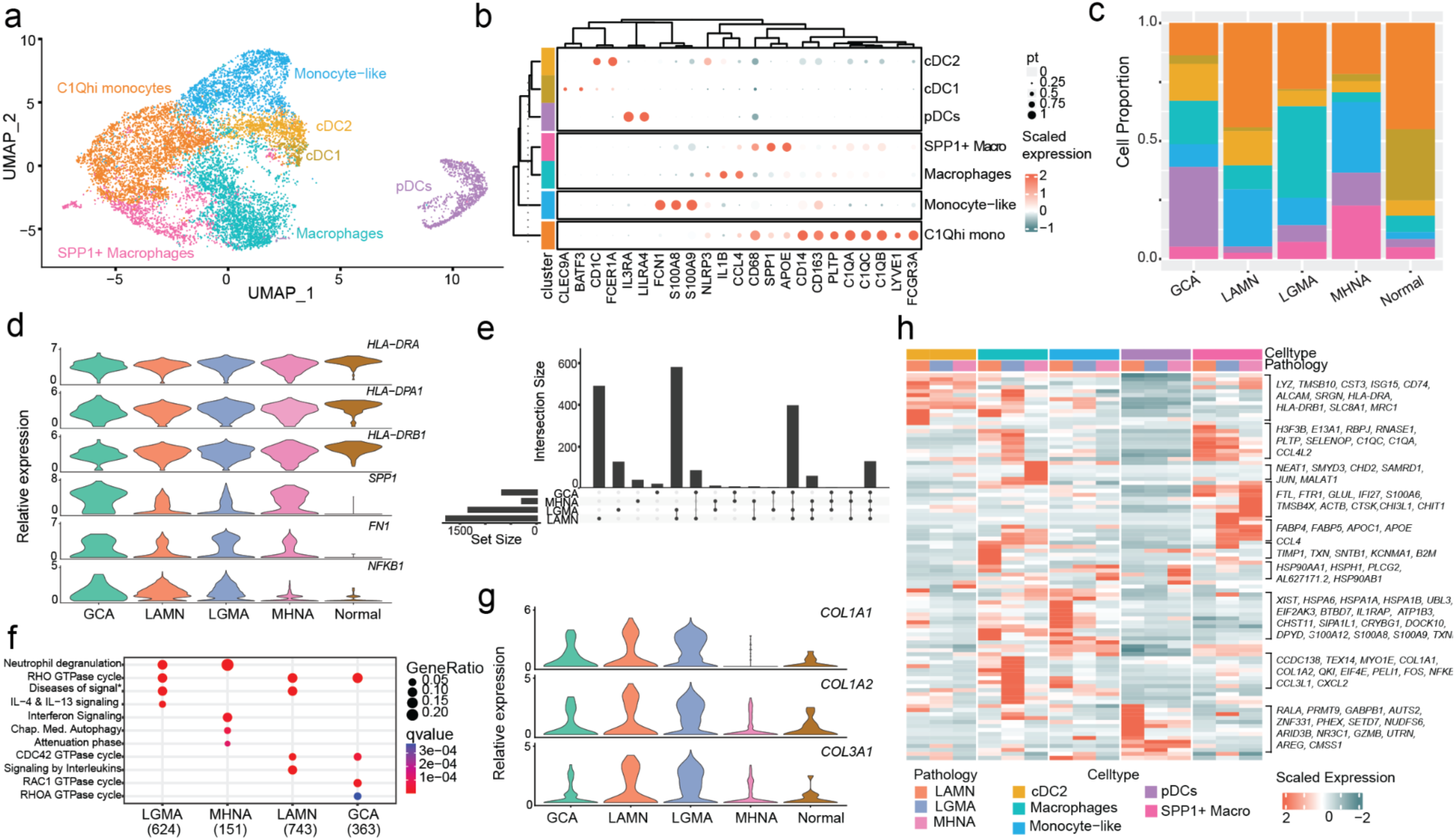
Characterization of the cancerous myeloid cell lineage. **a** UMAP plot displays different myeloid cells in all samples. **b** Signature marker gene expression of different myeloid subtypes. **c** Myeloid cell type proportion in different pathology groups and control. **d** C1Qhi monocytes in the cancerous samples express high levels of *SPP1*, *FN1* and *NFKB1* compared to normal cells. **e** Upset plot displays shared DEGs between each pathology vs. control cells in C1Qhi monocytes. **f** Dot plot displays the top four enriched pathways of C1Qhi monocytes in each pathology compared to control. **g** Violin plot shows an upregulation of collagen genes in macrophages in different pathologies. **h** Heatmap depicts the average expression of the top highly expressed genes in each pathology per cell type, the cell types used for this analysis have at least 40 cells per pathology per cell type.

C1Qhi monocytes were known to accumulate in Behçet’s disease and contribute to hyperinflammation^44^. Compared to controls, cancerous C1Qhi monocytes express high levels of myeloid tumor markers such as *SPP1, FN1* and the nuclear factor *NFKB1* (Fig. 4d). To study the differences among C1Qhi monocytes derived from different pathologic tissues, we compared gene expression in C1Qhi monocytes from each pathology group against normal cells (Fig. 4e). Upregulated signaling pathways in cancerous C1Qhi monocytes compared to control cells include IL-4 and IL-13 signaling (LGMA) and interferon signaling (MHNA) (Fig. 4f). Moreover, cancerous C1Qhi monocytes exhibit an M2-like phenotype with a significantly elevated expression of M2 signature genes compared to M1 signature genes across pathologies (Supplementary Fig. 11).

To further explore the molecular signatures of myeloid cells across pathologies besides C1Qhi monocytes, we examined gene expression patterns in the remaining myeloid cell types. For this analysis, we focused on cell types with at least 40 cells per pathology group. We identified pathology-specific gene expression profiles in the myeloid cells (cDC2, pDCs, Macrophages, Monocyte-like and *SPP1*+ macrophages) among LAMN, LGMA and MHNA pathologies (Fig. 4h). Interestingly, macrophages from cancerous tissues, especially from LGMA tissues, express high levels of collagen genes (*COL1A1, COL1A2* and *COL3A1*) (Fig. 4g-h), a phenotype previously reported in the airway macrophages of the fibrotic lungs^45^.

### Cancer-associated fibroblasts support fibrotic development in appendiceal cancer

The stromal compartment, which is made up of mesenchymal and endothelial cells, is crucial for tumor formation at all stages, from the start of aberrant proliferation to the emergence of therapeutic resistance. We observed higher cell numbers in the stromal compartment of cancerous tissues compared to control (Fig. 1f, Supplementary Table 3). The stromal compartment in our dataset consists of six mesenchymal cell types (Fig. 5a, Supplementary Fig. 12a-b) and two endothelial cell types (Supplementary Fig. 3). Normal mesenchymal cells consist of mostly fibroblasts, while myofibroblasts are present at a high number in the cancerous tissues and *FAP*+ cancer-associated fibroblasts (CAFs) are completely absent in the normal tissues (Fig. 5b, Supplementary Fig. 12c).

**Fig. 5:**
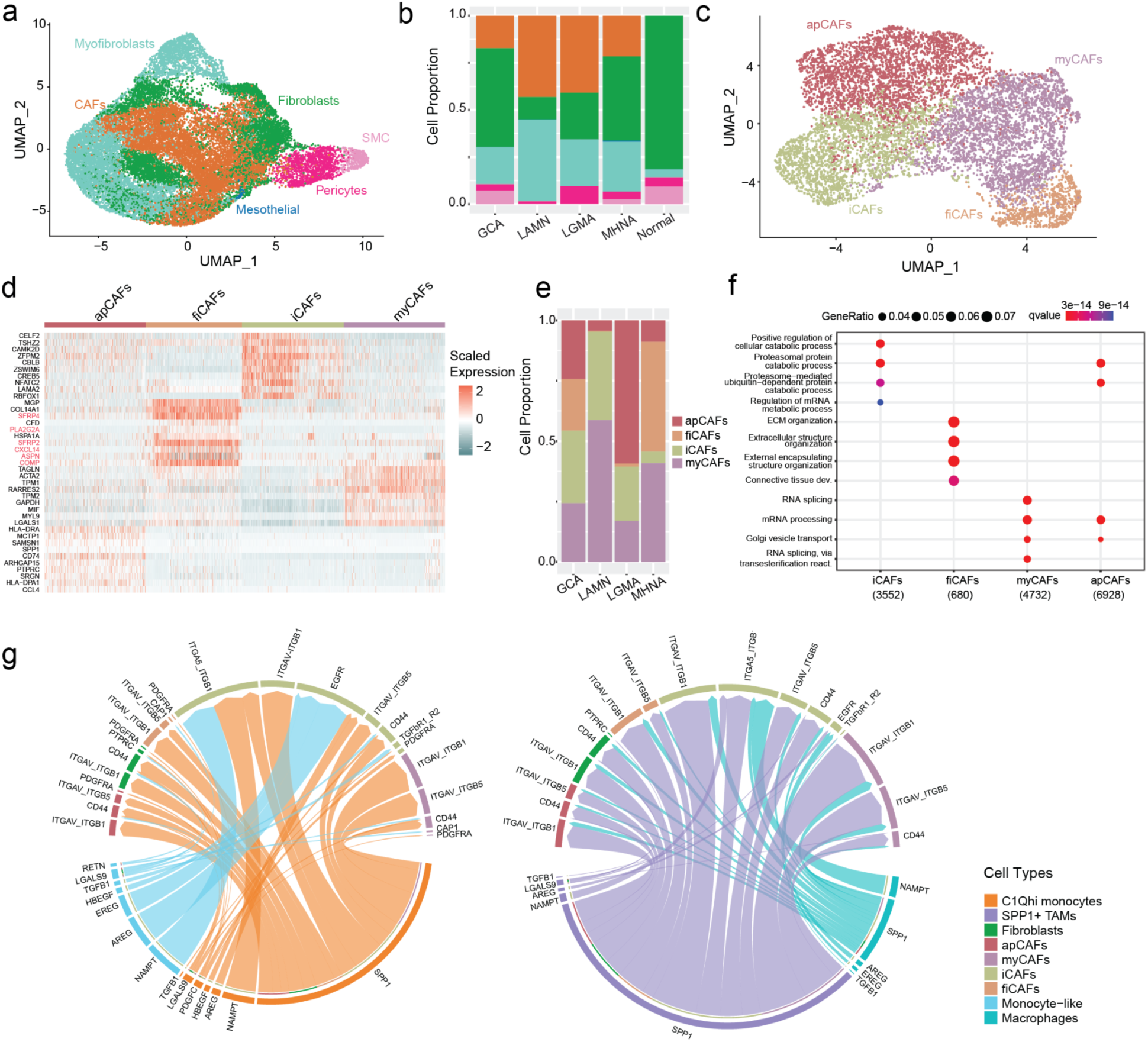
Characterization of mesenchymal cells in appendiceal cancer. **a** UMAP plot displays distribution of the six different cell types in the mesenchymal cell lineage. **b** Enrichment of CAFs in the appendiceal neoplasms samples. **c** UMAP plot displays distribution of the four different subtypes of CAFs. **d** Heatmap depicts relative expression (normalized and scale z-score) of the top ten upregulated genes in each CAF subtype, the genes known to be involved in fibrosis are highlighted in red. **e** Cell type proportion of each CAF subtype in each pathology. **f** Clusterprofiler GO enrichment analysis for genes upregulated in each CAF subtype (FDR ≤ 0.1). **g** Chord diagrams display receptor-ligand patterns and their weights in the interactions between the cancerous myeloid cells as senders (Left: C1Qhi monocytes and Monocyte-like; Right: Macrophages and *SPP1*+ macrophages) and cancerous fibroblast/CAF cells as receivers.

To further investigate the roles of CAFs in appendiceal neoplasms, we subsetted this cell population and re-clustered cells to group them into ten different clusters (Supplementary Fig. 12d). We then performed a DE analysis to examine the top genes uniquely expressed in each cluster, or shared among closely-related clusters, and identified four CAF subtype populations: inflammatory CAFs (iCAFs - clusters 2, 7, 8), antigen-presenting CAFs (apCAFs - clusters 0, 3), myofibrotic CAFs (myCAFs - clusters 1, 4, 5), fibrotic CAFs (fiCAFs - cluster 6) (Fig. 5c) and a small cluster (cluster 9, 33 cells) expressing mesothelial marker *UPK3B* (Supplementary Fig. 11d-e). Gene expression signatures used to identify CAF subtypes include chemokines *CXCL12/CXCL14* (iCAFs), myofibroblast markers *HOPX/MYL9* (myCAFs), CD74 and HLA-complex genes (apCAFs) and fibrotic genes (fiCAFs) (Supplementary Fig. 12e, Supplementary Fig. 13). fiCAFs express high levels of genes related to ECM organizations (Fig. 5f) and genes with known function in skin fibrosis (*SFRP2^46^*, *SFRP4^47^*, *COMP^48^),* liver fibrosis (*CXCL14^49^),* and lung fibrosis (*PLA2G2A^50^*, *ASPN^51^)* (Fig. 5d). Among CAFs subtypes, fiCAFs are detected predominantly in GCA and MHNA tissues (Fig. 5e), suggesting a role in higher grade appendiceal adenocarcinomas.

The interactions between CAFs and *SPP1*+ macrophages have been demonstrated to play pivotal roles in tumor growth and progression. To identify the key mediators of CAFs and *SPP1*+ macrophages interaction in appendiceal cancer, we explored cell-cell communication mechanisms of the four CAF subtypes and *SPP1+* macrophages (Fig. 5g). Among the different ligand-receptor interaction patterns detected, *SPP1*+ macrophages and other myeloid cells (macrophages, C1Qhi monocytes) interact with cancerous fibroblasts and CAFs through interactions of SPP1 and various integrins. Of note, iCAFs display the strongest interactions, and also a unique integrin alpha-5/beta-1 (ITGA5_ITGB1) - SPP1 interaction with C1Qhi monocytes and *SPP1*+ macrophages. In addition, iCAFs interact with monocyte-like cells through EREG/AREG and EGFR signaling, suggesting a role in promoting tumor progression.

### Pathologic differences in cell-cell interactions in appendiceal cancer

Recent research has highlighted the importance of cell-cell interactions in the tumor microenvironment (TME) for tumor growth and metastasis. To unravel the differences in cell-cell communication among different appendiceal pathologies, we evaluated the putative cell crosstalk using the R package MultiNichenet^52^. Given that MultiNicheNet infers group differences at the sample level for each cell type, it is critical to have enough cells per sample of each cell type for each group in the comparisons. Therefore, we combined some cell types into lineages to ensure there are enough cells (minimum of five) per sample and enough samples (minimum of two) per cell type for each pathology. This includes the tumor epithelial cells (goblet-like cells, SPINK4^hi^ cells and MUC5B^hi^ cells), endothelial cells (venous and arterial) and B cells (GALTB, GCB, Follicular B, Plasma B). In addition, since we focused on cellular crosstalk across the histology spectrum, we removed the control cells from the analysis.

The tumor cells communicate with each other and with other cell types in the TME to promote cancer progression and metastasis. Our analysis revealed pathology-specific cellular crosstalk within the tumor cells and between tumor cells and other cell types (Fig. 6, Supplementary Table 6). With the exception of LGMA, tumor cells from all pathologies display some levels of upregulated interaction within the tumor or with other cell types. GCA tumor cells (both as receivers and senders) exhibit the highest levels of cell-cell communication (Fig. 6a, 6e). Within GCA tumor cell population, we identified an enrichment in three ligand-receptor pairs: a phospholipase PLA2G2A and receptor tyrosine-protein kinase ERBB3, the heparin binding EGF-like growth factor (HBEGF) and CD9 antigen, or the paracrine hormone uroguanylin (GUCA2B) and the intestinal receptor Guanylyl cyclase C (GUCY2C). Among these interactions, HBEGF-CD9 is involved in gastric tumorigenesis^53^ and GUCY2C signaling plays essential roles in intestinal epithelial integrity^54^. Besides intracellular crosstalk, GCA tumor cells exhibit elevated communication with the tumor necrosis factor TNF from B cells and CD8+ T cells through TNFRSF1A receptor. As senders, GCA tumor cells express CD9 to interact with CD36 on the surface of C1Qhi monocytes and *SPP1*+ macrophages, an interaction known to participate in macrophage signaling in response to oxidized LDL^55^. Three ligand-receptor patterns are enriched for cellular crosstalk between tumor cells and iCAFs in GCA tissues: VEGFA-NRP1, known to enhance angiogenesis^56^, HBEGF-ERBB4, known to mediate HBEGF-induced metastasis activity,^57^ and ANG-PLXNB2 axis, known to regulate cancer cells^58^. Tumor cells in MHNA tissues have elevated interactions with C1Qhi monocytes through MMP9- EPHB2 ligand-receptor activity, an interaction that triggers cell-repulsive responses^59^. LAMN and LGMA tumor cells display an elevated interaction with iCAFs through the WNT1 inducible signaling pathway CCN4 and integrin ITGB1, with LAMN cells having a stronger signal. In LAMN tissues, we observed an increase in the melanoma inhibitory activity and integrins (MIA-ITGB1/ITGA5) signaling between tumor cells and fibroblasts.

**Fig. 6:**
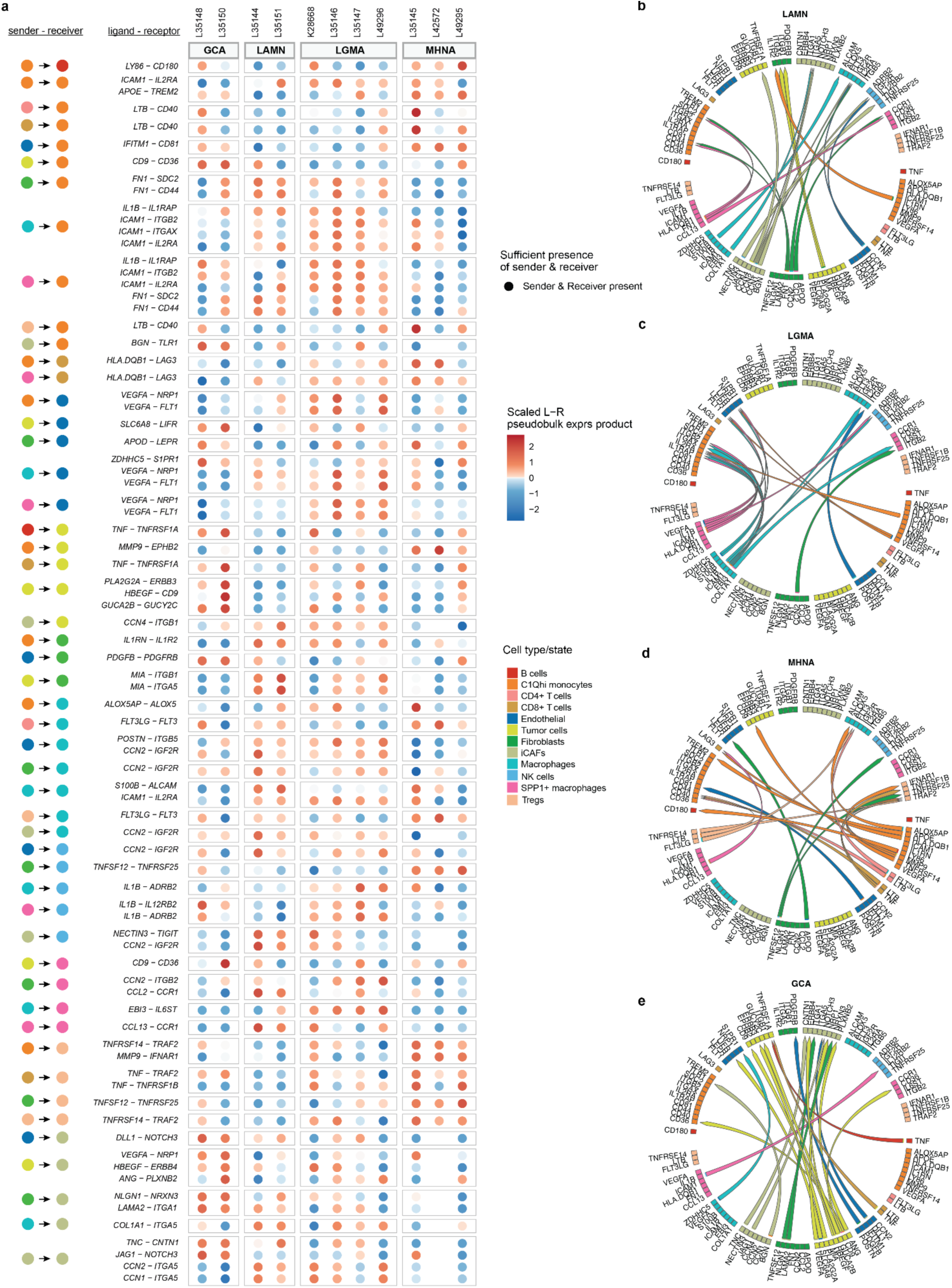
Cell-cell communication in the three appendiceal pathologies. **a** Heatmap depicts scaled pseudobulk ligand-receptor expression for the top 20 ligand-receptor pairs for each pathology, dot size illustrates whether a sample had enough cells (>= 5) for the cell type to be included in DE analysis. **b-e** Chord diagram depicts the top 20 ligand-receptor interactions in each pathology: LAMN (**b**), LGMA (**c**), MHNA (**d**) and GCA (**e**). The color of the arrow denotes the sender cell type that expresses the ligand, and the arrowhead denotes the direction from sender to receiver cell type.

The tumor cells are not the only cell type exhibiting differences in cellular crosstalk across the histologic spectrum. Differential cell-cell communication analysis with MultiNichenet revealed pathology-specific cell-cell communication patterns in many cell types in the dataset. For example, in MHNA, Tregs and C1Qhi monocytes have the highest number of cellular crosstalk with other cells, while iCAFs are predominant senders in LAMN and *SPP1*+ macrophages are more active in LGMA (Fig. 6b-d). Upregulation of specific ligands (LTB, FLT3LG) and receptors (TRAF2, IFNAR1, TNFRSF25, TNRFSF1B) on MHNA Tregs facilitate communication with many cell types, including C1Qhi monocytes, macrophages, CD8+ T cells and fibroblasts. Interestingly, the same sender-receiver pair may interact using different ligand-receptor patterns across pathologies. An example is the interaction between *SPP1*+ macrophages and C1Qhi monocytes: the ligand-receptor pairs IL1B-IL1RAP and ICAM1-ITGB2/IL2RA are enriched in LGMA, while the ligand-receptor pairs FN1-SDC2/CD44 are enriched in LAMN samples. Importantly, we identified inter-patient heterogeneity in cell-cell communication in many cell types across the histologic spectrum. Our analysis, therefore, provides an insight into the molecular differences across appendiceal neoplasm histology.

## Discussion

Although the integration of histologic findings and somatic mutation characterization have identified driver mutations (GNAS and KRAS) of appendiceal neoplasms, little is known about the central mechanisms underlying epithelial cellular remodeling and the tumor microenvironment. This study comprehensively explored the single-cell landscape of appendiceal neoplasms and characterized the molecular differences of cancer cell types/cell states across the histologic spectrum.

Goblet tumor cells have been proposed to contribute to the mucinous phenotype of appendiceal neoplasms^60^. We observed a similar phenotype with the tumor epithelial cells expressing high levels of goblet cell marker genes (*MUC2, TFF3, REG4* and *SPINK4*). An important observation from our study is that goblet-like cells derived from non-mucinous pathology (MHNA) express a different goblet marker (*TFF3*) compared to cells derived from mucinous pathology (LAMN, LGMA, GCA). This is consistent with previous studies in appendiceal neoplasms and CRC: high levels of *MUC2* were present in mucinous carcinoma but not in non-mucinous cancer^61,62^. Epithelial cells derived from GCA express high levels of *MUC5B*, a mucin gene expressed highly in adenomas and adenocarcinomas of the colon tissues^63^, and the transcription factor *SPDEF* (Supplementary Fig. 4c). Both *MUC5B* and *SPDEF* are among the signature genes driving invasive mucinous adenocarcinomas of the lung (IMA), and *SPDEF* induced *MUC5B* expression in human mucinous lung cancer cells^64^. The elevated expression of both *MUC5B* and *SPDEF* in GCA goblet-like cells suggests these genes may act similarly to IMA.

Immune cells, mainly T lymphocytes and myeloid cells, frequently infiltrate human tumors and are recruited to the tumor site by chemokines and cytokines secreted by various cells in the tumor microenvironment. While bulk RNA sequencing has suggested an immune-enriched subtype, the recruitment of immune cells in appendiceal neoplasms has not been characterized. We demonstrate here that there is significant heterogeneity of effector immune cell infiltration across the histologic spectrum with an abundance of CD8+ T cells in LGMA and LAMN samples in comparison to GCA and MHNA samples. Despite a significant shift in CD8+ T cells and NK cells frequency in tumor compared to control tissues, we were unable to detect significant enrichment of conventional immune checkpoint genes (*PD-1*, *CTLA4*) in tumor T lymphocytes, further confirming appendiceal neoplasms being immunologically cold and likely unresponsive to current anti-PD1/PDL1 or anti-CTLA4 checkpoint blockade^17^. However, the cancerous *CD4*+ *FOXP3*+ Tregs express high levels of antigen-presenting HLA class II (HLA-DR+, HLA-DP+ and HLA-DQ+) and tumor necrosis factor receptors (*TNFRSF4, TNFRSF9, TNFRSF14, TNFRSF18*). Tregs may play a dual role in cancer, either associated with poor prognosis in some cancers or favorable outcomes in others^65^. For example, *TNFRSF4-*expressing Tregs promote immune escape of chronic myeloid leukemia stem cells^66^ while *TNFRSF9* promotes immune suppressive activity in Tregs in lung cancer^67^. In addition, cell-cell communication analysis revealed that Tregs have the highest cellular crosstalk in MHNA tissues, mostly TNF signaling through TNFRSF23, TNFRSF1B or TRAF2 receptors. Taken together, tumor-infiltrated *CD4+ FOXP3+* Tregs may play an essential role in immune evasion of cancer cells in appendiceal neoplasms. The presence of effector immune infiltrates within appendiceal neoplasms presents an opportunity for the development of novel immunotherapies.

Other components of TME, myeloid and mesenchymal stromal cells, have been shown to form distinctive niches for tumor growth and metastasis^68^. It is known that *FAP*+ fibroblasts (CAFs) and tumor-associated macrophages (TAMs) maintain a bilateral interaction, in which M2-type macrophages can activate CAFs to promote tumor progression and CAFs can convert the M1 macrophages in TME into M2-like macrophages^69–71^. In this study, we identified a novel CAF subtype, fiCAFs, that express high levels of genes involved in fibrosis such as chemokine (*CXCL14*), *SFRP2, SFRP4,* and *ASPN*. fiCAFs are present at a higher frequency in MHNA and GCA, indicating the important role of fiCAFs in higher-grade appendiceal neoplasms. In appendiceal neoplasms, the interactions between myeloid cells (specifically C1Qhi monocytes and *SPP1*+ macrophages) and CAFs through SPP1-integrins ligand:receptor may play essential roles in TME supporting the tumor growth.

Cancer cells continuously communicate with each other and with other cell types in the TME to enable tumor growth and metastasis. Therefore, changes in cell-cell communication contribute significantly to tumor development. We identified significant differences in cellular crosstalk across the histologic spectrum, indicating different molecular mechanisms being involved in tumor growth and progression in each pathology. A key discovery from our analysis is the increase in tumor cell cellular crosstalk in more aggressive tumor (GCA) compared to moderate/high grade (MHNA) and low grade (LAMN, LGMA). The three ligand-receptor pairs being upregulated in GCA tumor intracellular crosstalk play essential roles in tumor initiation and progression (PLA2G2A-ERBB3^72^), tumorigenesis (HBEGF-CD9^53^) or maintaining the intestinal epithelial integrity (GUCA2B-GUCY2C^54^). GCA tumor cells send signals to communicate with C1Qhi monocytes and *SPP1*+ macrophages (CD9-CD36), endothelial (SLC6A8-LIFR) and iCAFs (VEGFA-NRP1, HBEGF-ERBB4, ANG-PLXNB2). Interactions between tumor cells and other cell types in the TME (myeloid cells, lymphocytes, stromal) are also different across pathologies and may have important impact on tumor development, metastasis and drug resistance^73^. Our cellular crosstalk analysis provides an insight into cell-cell communication in appendiceal neoplasms across the histologic spectrum and thus, will benefit therapeutic development.

Despite the insights uncovered by this study, there are many questions that remain unanswered. With the goal of delineating heterogeneity between histologic subtypes, we focused on parietal peritoneal metastases only. Our study does not characterize the cellular landscape of primary tumors, omental metastases, ovarian metastases or extra-peritoneal metastases. These sites likely harbor unique cell types and tumor microenvironments that warrant further study. Additionally, we have not performed characterization of peritoneal metastases from high-grade signet ring cell adenocarcinomas or high-grade mucinous adenocarcinomas; both of which are associated with poor prognosis. Lastly, while we ensured adequate sampling to characterize major cell types for histologies investigated, it is possible that rarer cell populations were not adequately represented. These limitations will be addressed in future studies.

In conclusion, our study provides, for the first time, high-resolution insights into the complexity and plasticity of appendiceal neoplasms across the histologic spectrum. We demonstrate that different histologies are composed of unique cancer epithelial lineages with distinct signaling mechanisms. Similarly, cancer cells orchestrate a unique tumor microenvironment that substantially differs across the histologic spectrum. Despite the low expression of conventional immune checkpoint markers, we unexpectedly identified an abundance of effector immune subsets to varying degrees. By uncovering cancer, stromal, and immune cell circuitries, our findings open a plethora of opportunities for histology-specific therapeutic development.

## Materials and Methods

### Sample collection and preparation

This study was approved by the City of Hope Comprehensive Cancer Center, Institutional Review Board (IRB: 18209). All study subjects in this study provided written informed consent for sample collection and data analyses. Cells were isolated and dissected from appendiceal neoplasms. Once trimmed, tissue samples were stored in cold RMPI medium, then quickly was cut to small pieces (1-3 mm) in petri dish with 3 -5 ml RMPI medium containing 0.05mg/ml TM Liberase (Sigma Aldrich, 5401119001) and 0.02 mg/ml DNAaseI (ThermoFisher, EN0521). Minced tissue was transferred to 50 ml conical tubes, digested at 37 ° C for 30 mins with rotation at 150 rpm, and then gently pipetting with 5 ml serological for 25-30 mins, followed by adding pre-warmed full RMPI medium (supplemented with 10% FBS) to quench the digestion. Cells were filtered through a 70 µM and 37 µM cell strainer to isolate single cells. The cell suspension was centrifuged at 500xg for 10 mins and resuspended with 5 ml ACK lysis buffer (K.D Medical, RGF3015) at room temperature for 3 mins to remove red blood cells. The cells were washed with 5 ml warmed full RPMI. Microbeads (Miltenyi Biotec, 130-090-101) were used for dead cell removal. The cells were washed twice and resuspended in DPBS (Sigma Aldrich, D4537) with 0.04% Bovine Serum Albumin. Cell viability and number were detected using a TC Automated Cell Counter (Bio-Rad Laboratories), and samples with at least 70% cell viability were processed.

### Histology and pathology classification of samples

The patients were classified based on the PSOGI and WHO consensus definitions for appendiceal cancer^8, 9^. The distinction between peritoneal metastases from LAMN or LGMA was based on the primary tumor characteristics. Two expert pathologists - LA and RM - independently reviewed and classified patient specimens. Peritoneal deposits were classified in correlation with the primary tumor in the appendix. Only samples with unambiguous histologic classification were selected for single cell sequencing and analysis.

### Identification of somatic mutations in appendiceal neoplasms samples

Somatic mutation identification was performed using GEM ExTra test, a clinical test performed by Ashion Analytics (now part of Exact Sciences Corp.), using standard protocol on FFPE tumor sections^21^.

### scRNA-seq library preparation and sequencing

Cells were captured on a Chromium controller (10X Genomics) using a single cell 3’ Reagent kit V3.1 (10X Genomics, PN-1000121) targeting ∼5000-10000 cell/sample. Single cell RNA-seq libraries were prepared by following the manufacturer’s instruction. cDNA and sequencing libraries were analyzed on a High Sensitivity DNA Chip (Agilent Technologies, #5067-4626) to determine the cDNA and library quality. The library concentration was determined with a Qubit High Sensitivity DNA assay Kit (ThermoFisher Scientific). The libraries were sequenced on Illumina NovaSeq 6000 platform (Illumina) at The Translational Genomics Research Institute (TGen) with the sequencing depth of 50K-100K reads/cell with 1% phiX using the following cycles: Read 1 (101 cycles), i7 (8 cycles) and Read 2 (101 cycles). Reads with a read quality of less than 30 were filtered out and CellRanger Count v5.0.2 (10X Genomics) was used to align reads onto the GRCh38 (Ensembl 98) reference genome.

### scRNA-seq data processing

Seurat v4.3.0^74^ was used to perform data integration, dimensionality reduction, unsupervised clustering, and data analysis for the scRNA-seq data. Individual CellRanger output files were read into Seurat v4.3.0 to generate a unique molecular identifier count matrix that was used to create a Seurat object for each individual, and the percentage of reads mapped to mitochondria was calculated for each sample. Cells containing less than 500 identified genes or more than 25% of reads from mitochondria were eliminated from the object to remove low-quality cells. Ambient RNA removal for individual Seurat objects was identified using SoupX^75^ and doublet cells were identified using DoubletFinder^76^ with a 5% doublet formation rate and were removed from downstream analysis. Seurat fast integration using the reciprocal PCA (RPCA) pipeline was used to integrate all samples from each sequencing batch to eliminate batch effects. FindPC^77^ was used to select the optimal number of principal components (PCs) for UMAP visualization.

### Cell type annotation

cell type annotation was performed at two levels. For the first level, markers specific for each cell population were used to identify different cell lineages: *EPCAM+* (epithelial cells), *CD3E+* (T cells), *CD68+* (myeloid cells), *CD79A+* (B cells), *CXCR2+ FCGR3B+* (*FCGR3B+* neutrophils), *DCN+ THY1+* (mesenchymal cells) and *PECAM1+* (endothelial cells) (Supplementary Fig. 1b). Then, each cell lineage was subsetted from the combined object. New variable genes for each lineage were identified using the Seurat FindVariableGenes function on the raw RNA counts. Seurat ScaleData was used to re-scale the integrated assay using these new variable genes. Each cell lineage object then underwent dimensionality reduction and cluster splitting using Seurat RunUMAP, FindNeighbors, and FindClusters functions. Cell type annotation was performed on each lineage using a combination of cell type-specific markers (Supplementary Fig. S2b) and cluster-specific genes identified by the Seurat FindAllMarkers function on each cluster (please see supplementary figures for more information). Additional doublets were identified at this step as cells expressing markers for different cell types and were removed from the analysis. The final cell type annotation was projected back on the original integrated object containing all cells.

### Copy number variant (CNV) analysis

Infercnv (inferCNV of the Trinity CTAT Project. https://github.com/broadinstitute/inferCNV) was used to identify gains or deletions of whole chromosomes or large segments of chromosomes. Briefly, raw gene data were extracted from the Seurat object following infercnv “Using 10X data” section. The infercnv object was generated using the function CreateInfercnvObject with cells derived from normal appendix samples used as reference. Next, infercnv::run function was used to perform infercnv operations using a cutoff level of 0.1 (recommended by infercnv for 10X Genomics data), analysis_mode being “subclusters” to partition cells into groups with consistent patterns of CNV using Hidden Markov Model (HMM_type=i3 to predict the three-state CNV: deletion, neutral and duplication), “ward.D2” as clustering method and “random_trees” as the method used for partitioning the hierarchical clustering tree. A total of 10,644 genes were used in infercnv analysis on the epithelial cell population (5,205 cells with 4,763 tumor cells). The top 10 HMM features were added to the Seurat object using the function infercnv::add_to_seurat, and the proportion of expressed genes part of a CNV in a given chromosome per cell was plotted on the UMAP space using the Seurat function FeaturePlot.

### Differential expression analysis

prior to differential expression (DE) analyses, the Seurat function PrepSCTFindMarkers function was performed to correct the SCT counts due to the different SCT models used for the integrated pipeline. DE analyses were then performed using the Seurat FindMarkers function on the SCT raw count data matrix using the negative binomial test with Flowcell as a covariate and the adjusted p-values were calculated using “Bonferonni” correction.

### Studying cell-cell communication network

CellChat R package^78^ was used to perform receptor-ligand analysis for fibroblasts, CAFs and myeloid cells in the cancerous tissues (https://github.com/sqjin/CellChat). Specifically, we subsetted the cancerous cells for the myeloid cells (macrophages, SPP1+ macrophages, C1Qhi monocytes and monocyte-like), fibroblasts and CAFs. Then, cell-cell communication analysis was performed using CellChat ligand-receptor “secreted signaling” database. CellChat function *createCellChat* was used to create a CellChat object and cell types with less than 40 cells per group (in at least one pathology group) were removed from the analysis. All the steps including computing communication probability and inferring cellular communication network, inferring the cell-cell communication at a signaling pathway level and calculating the aggregated cell-cell communication network were performed using CellChat default parameters. Ligand-receptor patterns were visualized using the CellChat function *netVisual_chord_gene* using the myeloid cells as senders and fibroblasts/CAFs cells as receivers.

### Differential cellular crosstalk analysis

Multinichenet R package^52^ was used to study the differences in ligand-receptor interactions among pathology groups. For this analysis, we used a cut-off of at least two samples with 5 cells per cell type per comparison. We combined cell types into cell lineages for the endothelial, epithelial and B cells since some of the cell types from these lineages are present at a low number in one of the comparisons. Due to a low number of cells in many cell types in the normal appendix and goblet-cell adenocarcinoma samples, we removed these from the analysis. The Multinichenet analysis in this paper was performed following the guidelines from https://github.com/saeyslab/multinichenetr for a three-wise comparison analysis using the Nichenet v2 ligand network and ligand-target matrices. We adjusted the contrasts_oi argument from the three-wise comparison vignette for our comparison: "’LAMN-(LGMA+MHNA+GCA)/3’,’LGMA-(LAMN+MHNA+GCA)/3’,’MHNA-(LGMA+LAMN+GCA)/3’,’GCA-(LGMA+LAMN+MHNA)/3’". We included “Flowcell” as a covariate in the Multinichenet analysis to account for batch effects caused by different sequencing batches. Multinichenet analysis was performed using the recommended criteria to prioritize ligand-receptor interactions from the vignette and the top 20 ligand-receptor predictions for each pathology (GCA, LAMN, LGMA or MHNA) were chosen for visualization in Fig. 6.

### Pathway enrichment analysis

ClusterProfiler R package^79^ was used to perform pathway enrichment analysis. Specifically, significant DEGs (FDR ≤ 0.05 & avg_log2FC ≥ 2) were selected for each group, and the function compareCluster was used to identify enriched pathways. Dotplot was generated using the top 4 enriched pathways per group.

### Signature gene set score analysis

To calculate the signature gene set scores, we used the Seurat AddModuleScore function, with signature gene sets for M1 (*IL1A, IL1B, IL6, NOS2, TLR2, TLR4, CD80, CD86*) and M2 (*CD115 (CSF1R), CD206 (MRC1), PPARG, ARG1, CD163, CD301 (CLEC10A), Dectin-1 (CLEC7A), PDL2 (PDCD1LG2)* and *FIZZ1 (RETNLB)*). Boxplot was generated using ggpubr package and statistical analysis was performed using the Student T-test.

## Data and code availability

Raw and processed 10X genomics data generated in this study are available at GEO with the accession number GSE244031. All codes used to analyze data are available at https://github.com/tgen/banovichlab/tree/master/Appendiceal_neoplasms

## Acknowledgments

The authors acknowledge funding to M.R. and N.E.B. from Kure-It Cancer Foundation; as well as funding from an anonymous donor to MR that supported this work.

## Author contribution

L.T.B., N.E.B., and M.R. conceived and designed the analysis. Sample collection was performed by M.R., K.I., Y.W., and processing was performed by X.C., J.W., F.M. Appendiceal histology classification was performed by L.A. and R.M. L.T.B., M.R. and N.E.B. performed quality checks, data integration, and computational analyses. L.T.B., N.E.B., and M.R. analyzed and interpreted scRNA-seq data. L.T.B., N.E.B., and M.R. wrote and revised the manuscript, with significant input from K.I. All authors read and approved the manuscript before submission.

## SUPPLEMENTARY INFORMATION

**Supplementary Table 1.** Sample demographics information.

|  | Control | Cancerous |
| --- | --- | --- |
|  | n=5 | n=11 |
| Age | 52.4 [36-58] | 61.82 [39-74] |
| Sex (male) | 0 | 5 (45.46%) |
| Ancestry |  |  |
| Non-Hispanic white | 3 (60%) | 9 (81.82%) |
| Hispanic white | 2 (40%) | 1 (9.09%) |
| African-American | 0 | 1 (9.09%) |
| Pathology |  |  |
| Low-grade appendiceal mucinous neoplasm |  | 2 (18.18%) |
| Low-grade mucinous adenocarcinoma |  | 4 (36.36%) |
| Mod/High-grade non-mucinous adenocarcinoma |  | 3 (27.27%) |
| Goblet-cell adenocarcinoma |  | 2 (18.18%) |

**Supplementary Table 2: Detailed information about pathology classification and mutation data for all samples**

**Supplementary Table 3: Number of cells per cell type per diagnosis (csv files)**

**Supplementary Table 4: Cell proportion Anova test from Propeller (excel file)**

**Supplementary Table 5: Differential expression analysis (zip file)**

**Supplementary Table 6: Ligand-receptor prioritization table, full data from MultiNichenet**

---

**Supplementary Fig. 1:**
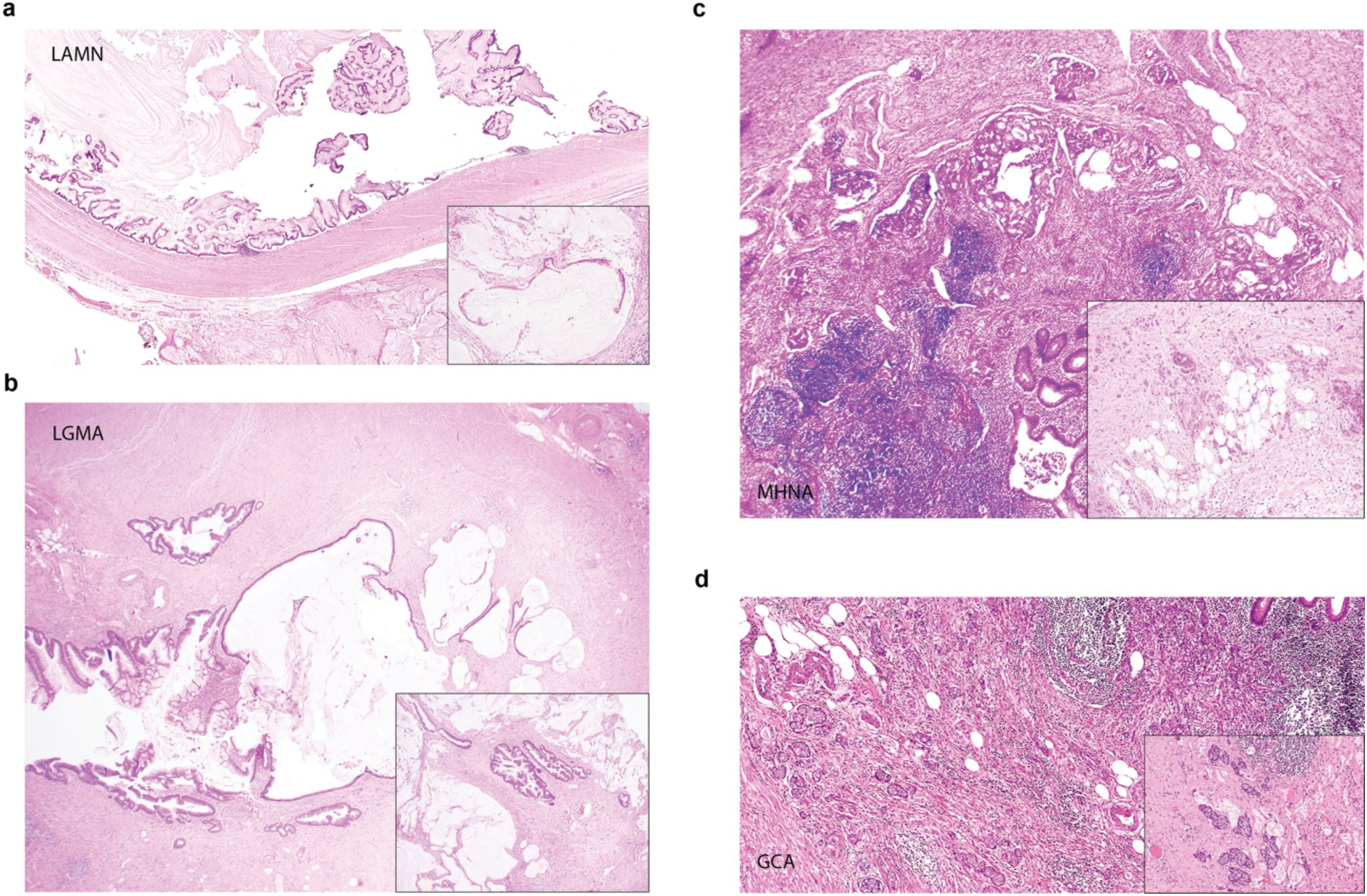
Pathology classification of appendiceal neoplasms in the study. **a** *Low grade appendiceal mucinous neoplasm (LAMN):* Appendix with low grade appendiceal mucinous neoplasm and periappendiceal mucin: inset shows peritoneal involvement with floating low grade mucnous neoplastic epithelium (H&E x10). **b** *Low-grade mucinous adenocarcinoma (LGMA)*: Appendix with involvement by low grade mucinous adenocarcinoma; inset shows infiltrating glands into the peritoneal soft tissue (H&E x 10). **c** *Moderate / high grade non-mucinous adenocarcinoma (MHNA)*: Moderately differentiated non-mucinous adenocarcinoma involving appendix; inset shows peritoneal deposit by high grade carcinoma (H&E x 10). **d** *Goblet cell adenocarcinoma (GCA):* Appendix with involvement by goblet cell adenocarcinoma; inset shows peritoneal involvement by goblet cell adenocarcinoma (H&E x 10).

**Supplementary Fig. 2:**
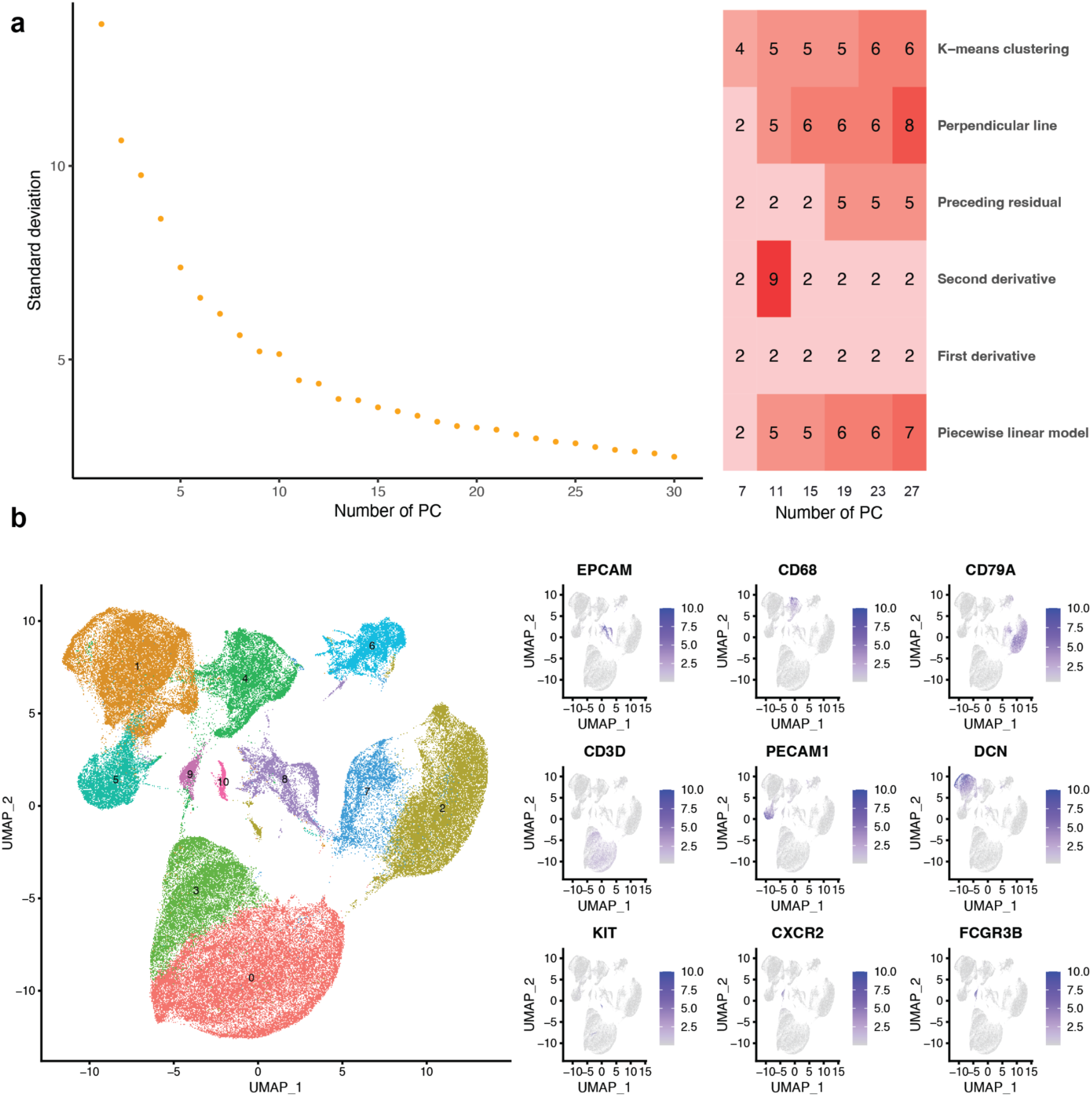
Selection of the number of principal components for dimensionality reduction in the appendiceal data set and cell lineage annotation. **a** The number of PCs used for dimensional reduction was determined based on the elbow plot and six different methods were calculated based on the elbow plot by findPC package, for this dataset, we used 15 PCs for dimensional reduction. **b** Expression of cell lineage-specific markers was used to assign cell clusters into cell lineages.

**Supplementary Fig. 3:**
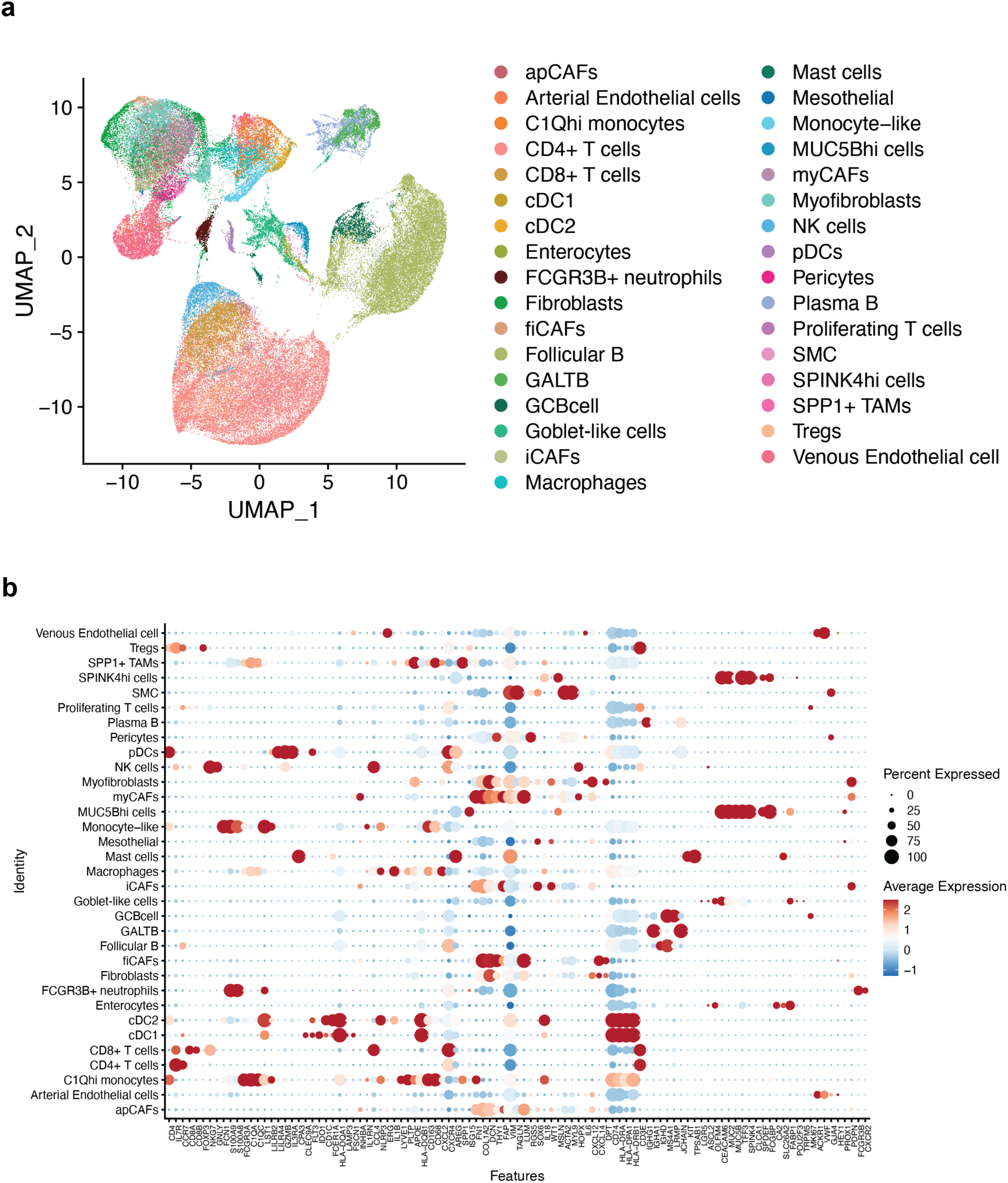
Granular cell type annotation in the appendiceal scRNA-seq dataset. **a** UMAP displays 32 distinct cell types in the scRNAseq dataset. **b** Expression of molecular markers used for granular cell type annotation.

**Supplementary Fig. 4:**
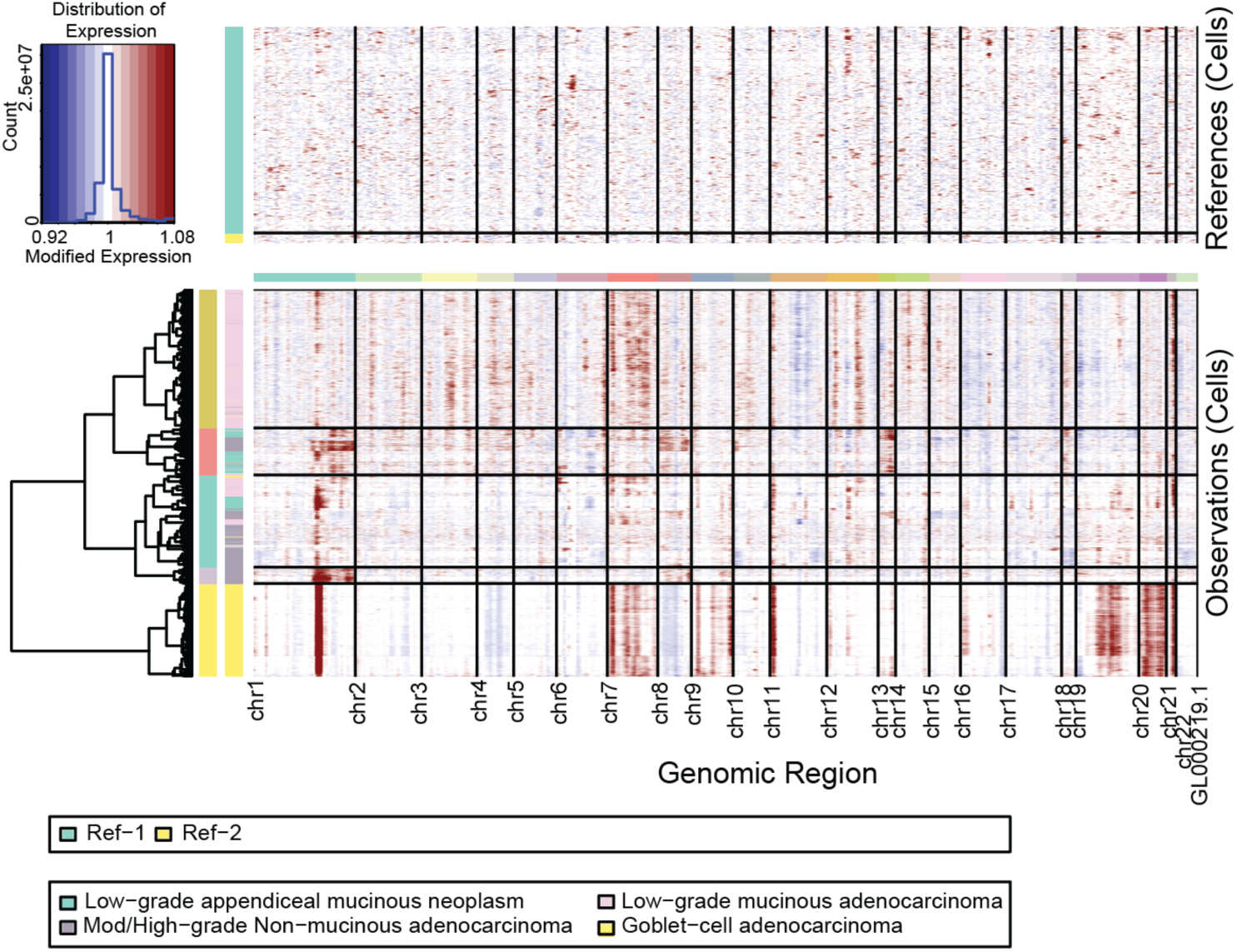
Heatmap displays CNVs analysis of the epithelial cell lineage, with rows corresponding to the individual cells and columns displaying genes. Heatmap color intensities represent the residual expression values, chromosomal duplication regions are in red blocks while chromosomal deletion regions are in blue blocks. The normal cells (Reference) at the top of the heatmap define baseline expression for genes in normal cells. The appendiceal tumor cells (Observations) are clustered based on CNVs profile into five distinct subgroups.

**Supplementary Fig. 5:**
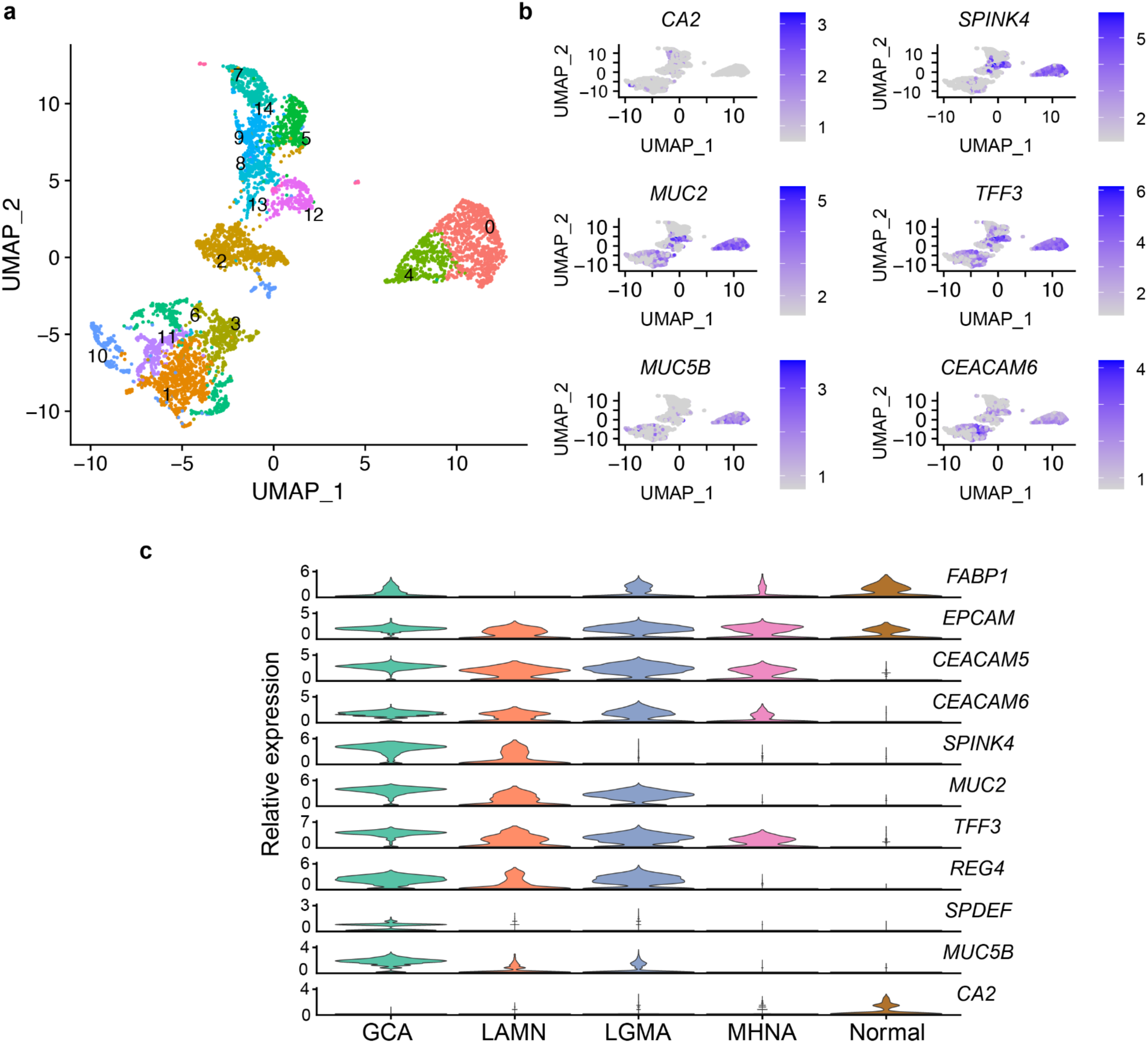
**a** UMAP plot displays cell clusters in the epithelial cell lineage. **b** Expression of cell type-specific marker genes in the epithelial cell lineage. **c** Violin plot depicts relative gene expression of marker genes in the epithelial cells across pathologies.

**Supplementary Fig. 6:**
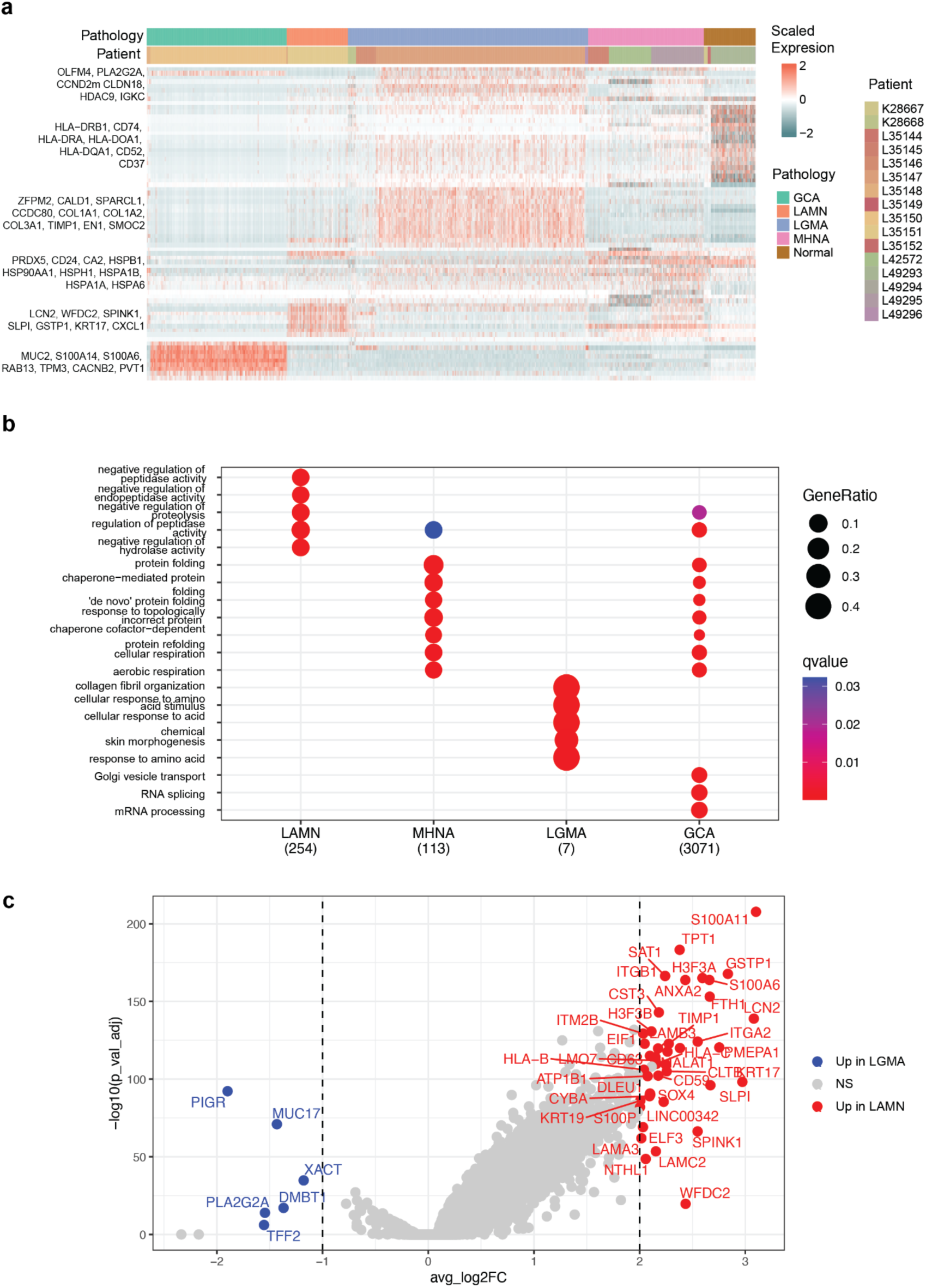
**a** Heatmap depicts the top DEGs in each pathology in the epithelial cell lineage. **b** Top 5 enriched GO terms in the highly expressed genes (FDR < 0.1 & avg_log2FC ≥ 1) in epithelial cells derived from different pathologies. **c** Volcano plot depicts DEGs in the goblet-like cells between LAMN and LGMA, labeled are significant DEGs (FDR ≤ 0.05, avg_log2FC ≥ 2 or < -1)

**Supplementary Fig. 7:**
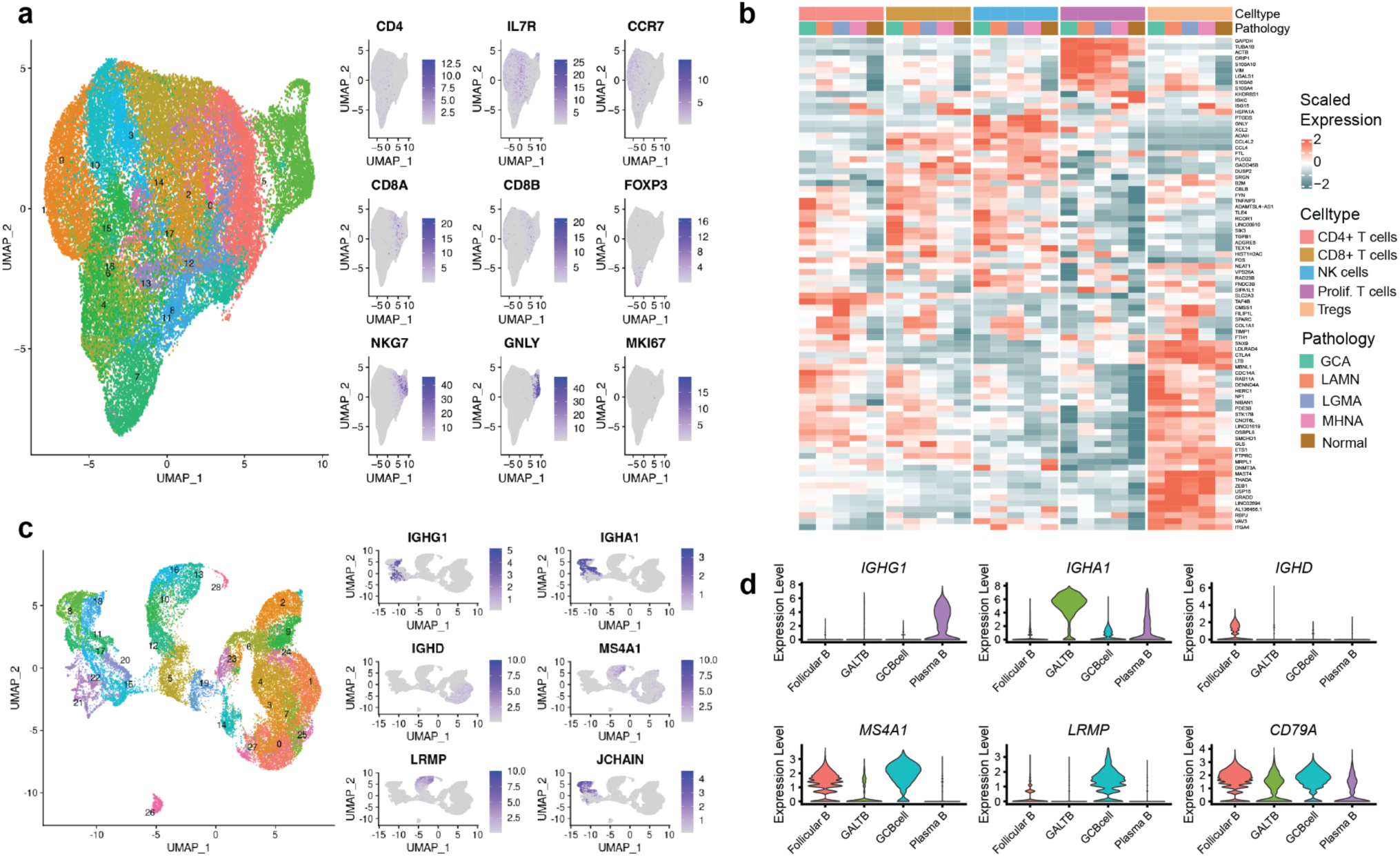
Cell type annotation in the lymphoid cell lineage. **a** UMAP plot depicts different clusters and expressions of canonical markers in the T cell population. **b** Heatmap depicts scaled expression levels of the top DEGs in each cell type per pathology group. **c** UMAP displays different clusters and expressions of canonical markers in the B cell population. **d** Violin plot depicts the expression of different cell type-specific markers in the B cell population.

**Supplementary Fig. 8:**
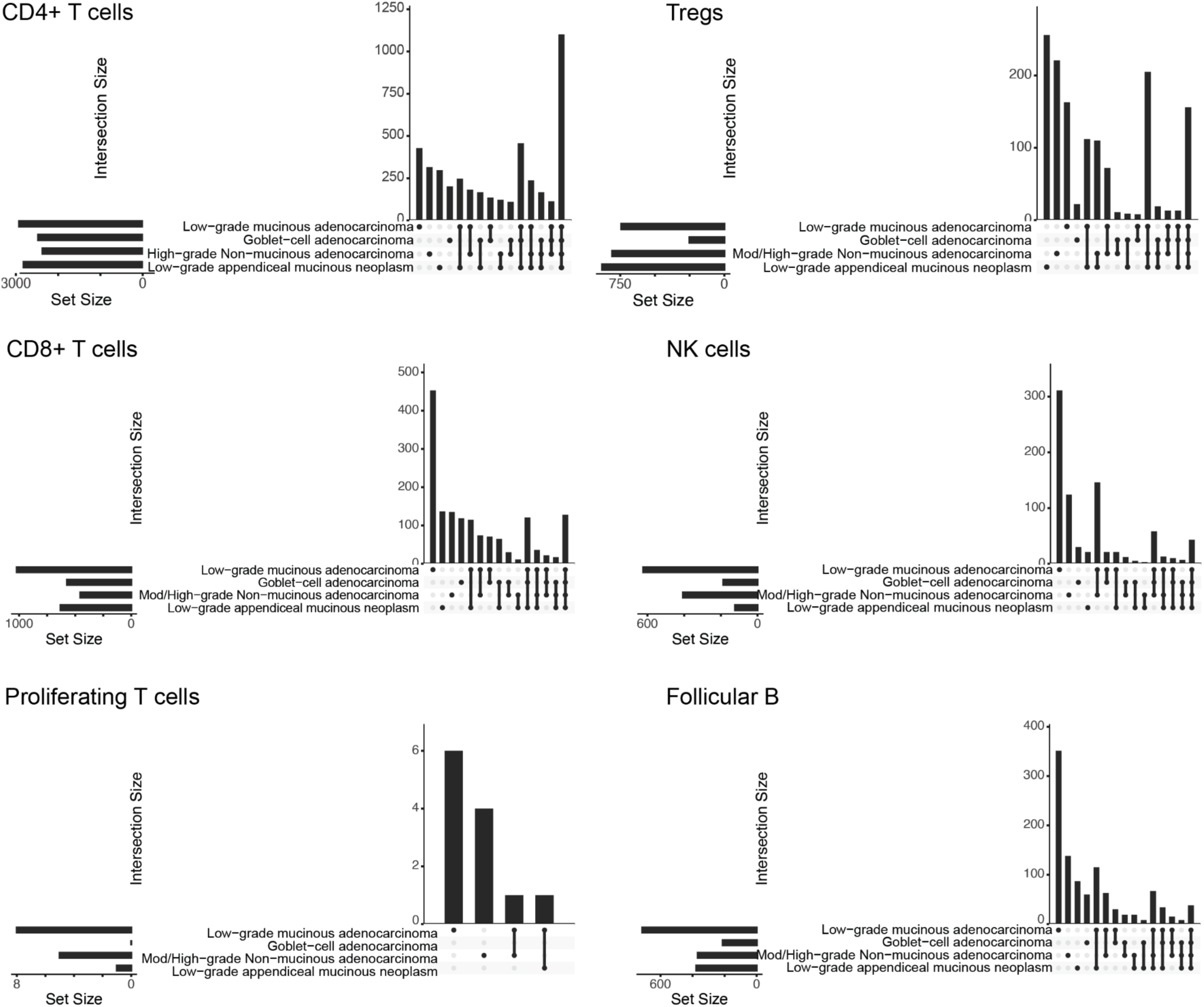
Upset plots display shared DEGs between different pathologies vs. control cells in the lymphoid cell lineage.

**Supplementary Fig. 9:**
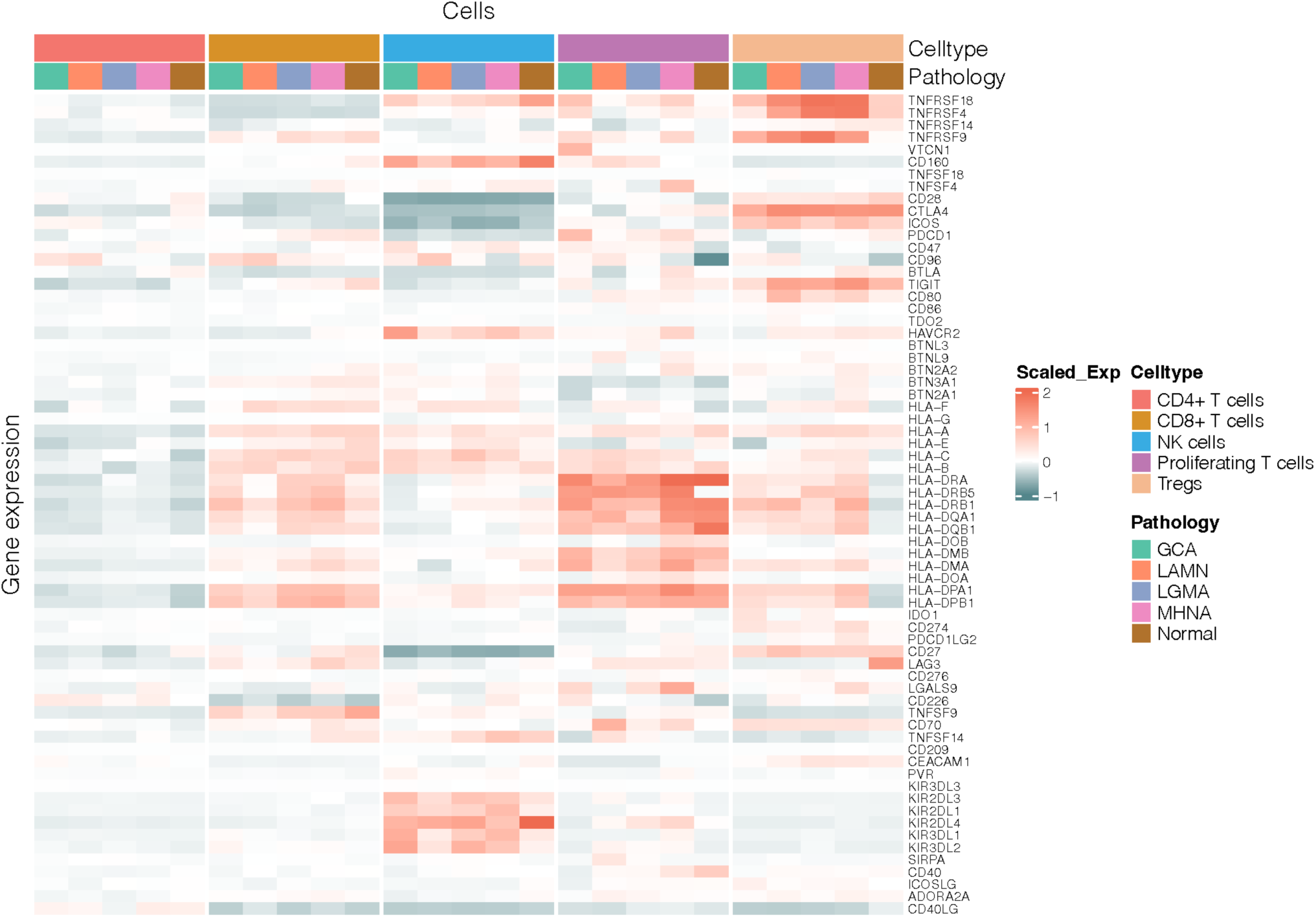
Heatmap depicts the average expression of immune checkpoint genes (ICGs) in different cell types of the T cell lineage. The list of ICGs used in this heatmap was adapted from ^80^.

**Supplementary Fig. 10:**
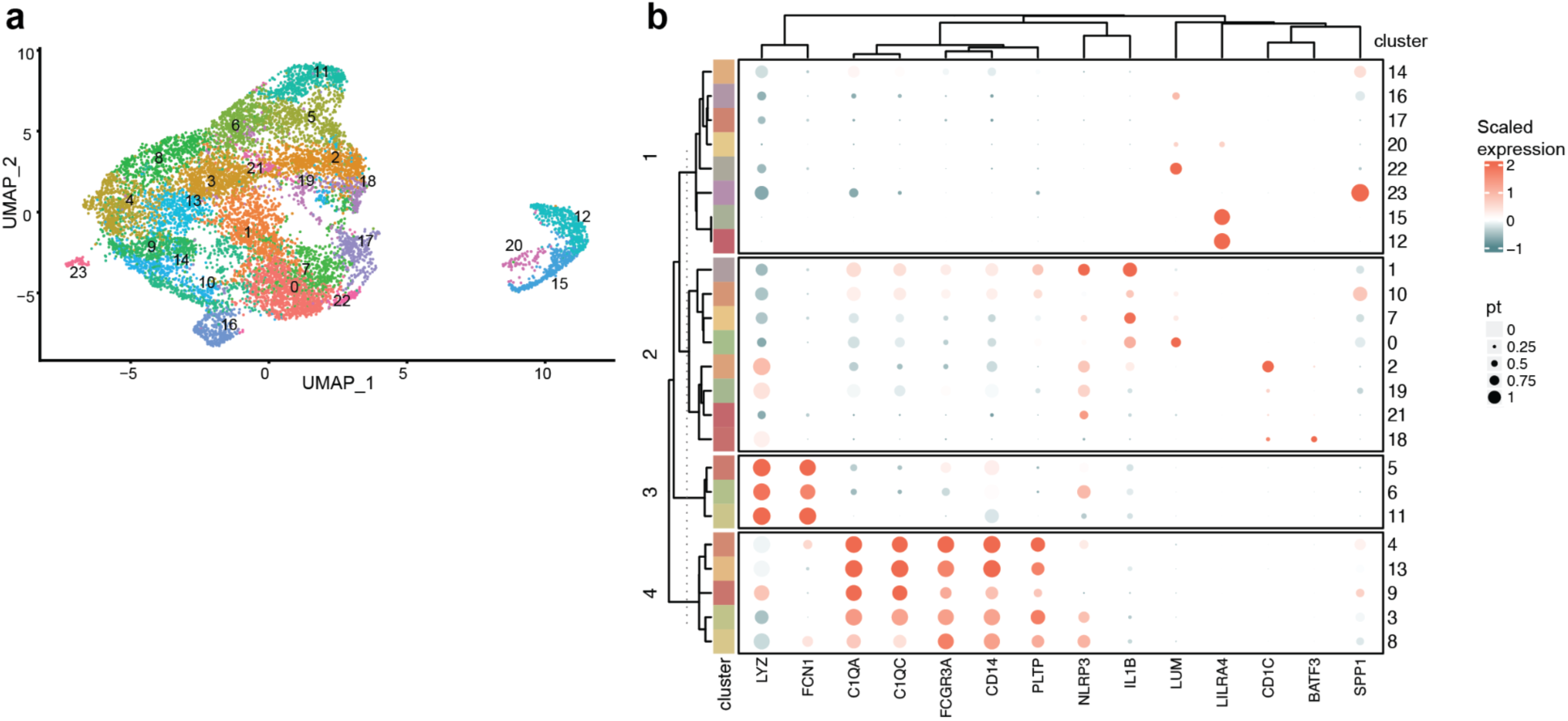
Cell type annotation in myeloid cell lineage. **a** UMAP plot depicts unsupervised cell clusters in the myeloid cell lineage. **b** Dotplot heatmap depicts expression levels of the myeloid cell markers for each cluster.

**Supplementary Fig. 11:**
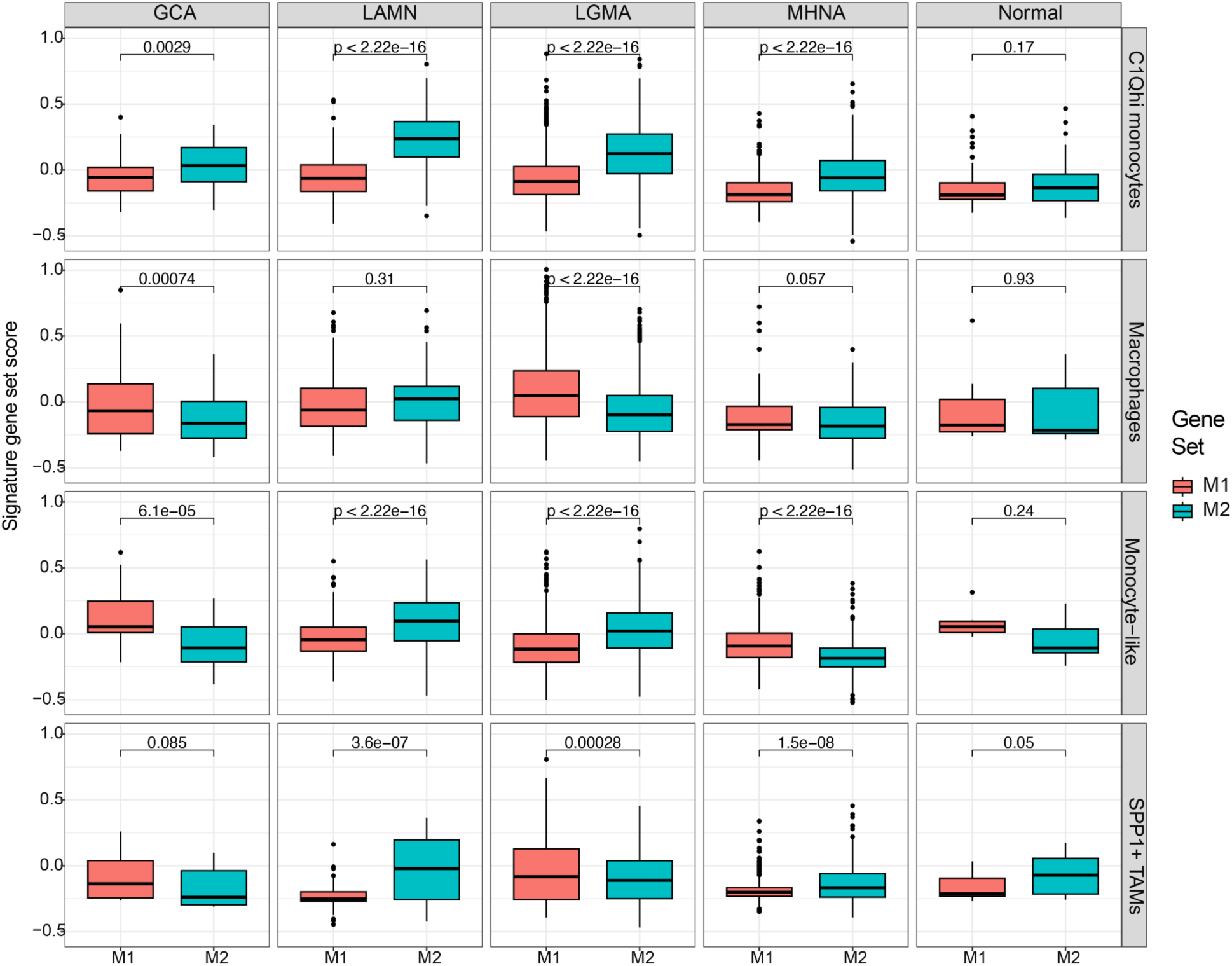
Signature gene scores for M1 or M2 macrophages in different myeloid cell types per diagnosis. The statistic test was performed using ggpubr stat_compare_means with Student T-test.

**Supplementary Fig. 12:**
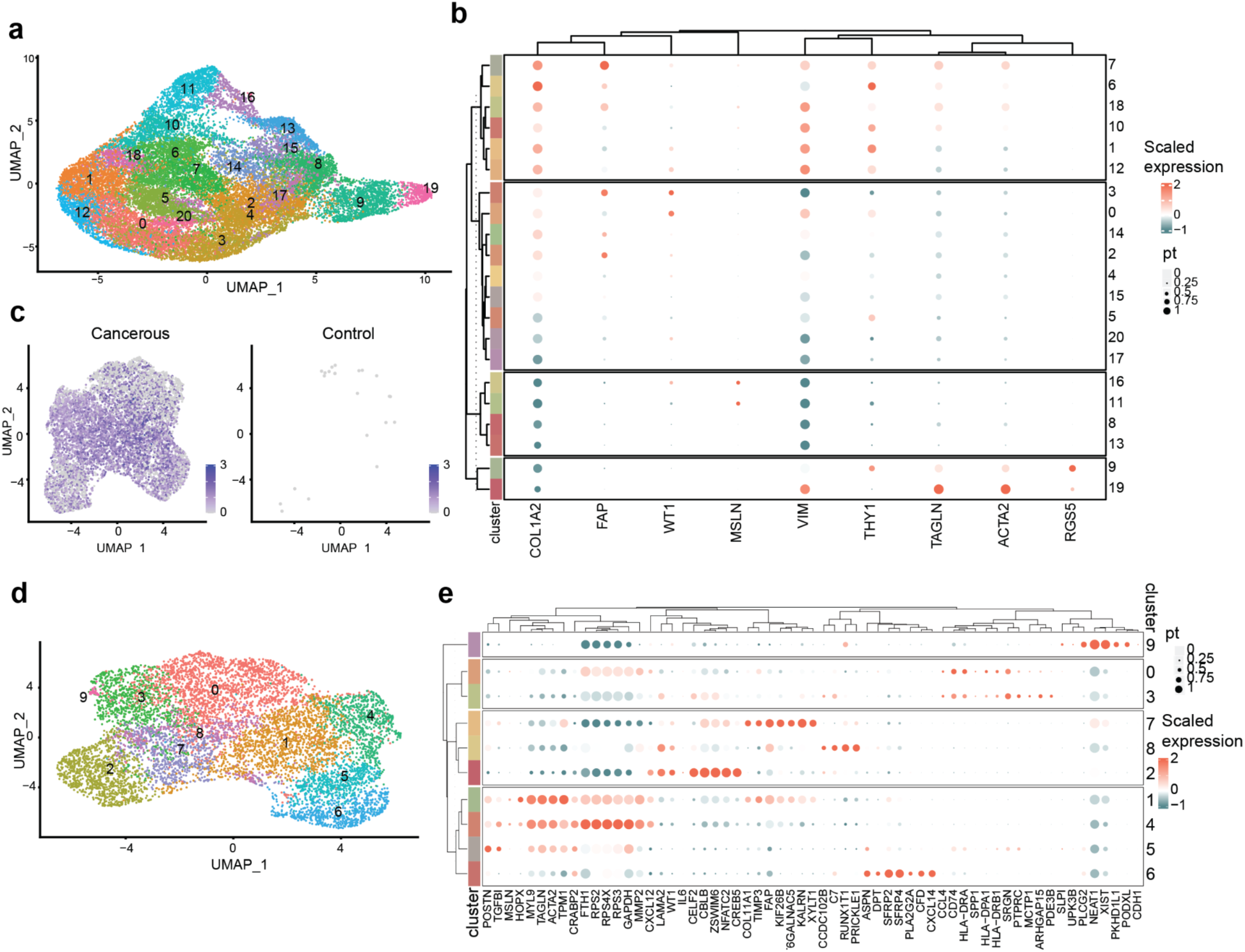
Cell type annotation in the mesenchymal cell lineage. **a** UMAP plot displays different cell clusters in the mesenchymal cell population. **b** Dotplot shows unique marker gene expression in each cell cluster. **c** No expression of *FAP* marker was detected in the control cells grouped together with CAFs, the scale bar displays scaled_expression level. **d** UMAP plot displays cell clusters in the CAFs population. **e** Dotplot displays marker gene expression in different CAFs cell clusters.

**Supplementary Fig. 13:**
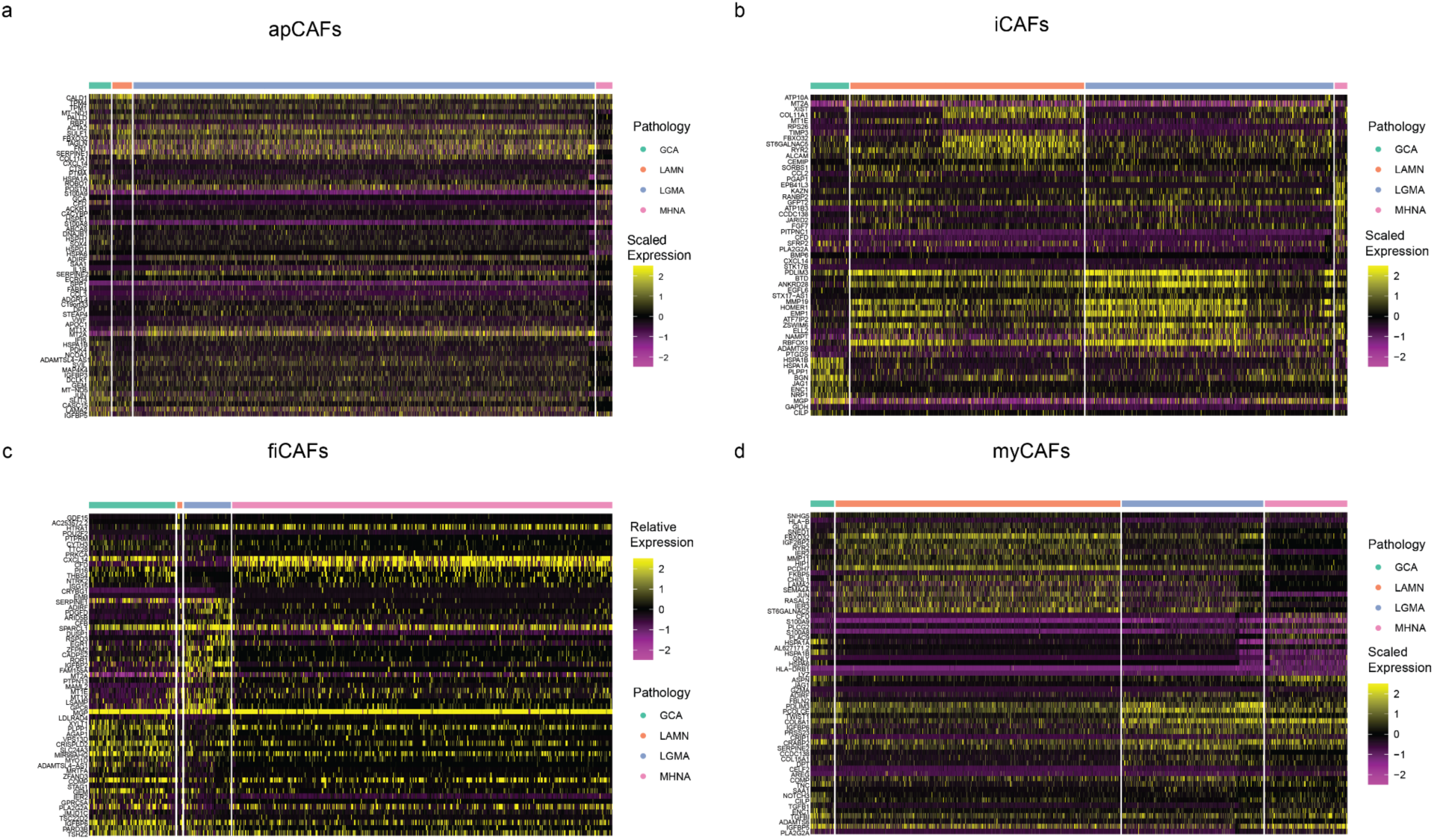
Heatmap depicts differences in gene expression (normalized and scaled) levels across pathology groups in each CAF subtype.

